# VariantFormer: A hierarchical transformer integrating DNA sequences with genetic variations and regulatory landscapes for personalized gene expression prediction

**DOI:** 10.1101/2025.10.31.685862

**Authors:** Sayan Ghosal, Youssef Barhomi, Tejaswini Ganapathi, Amy Krystosik, Lakshmi Krishnan, Sashidhar Guntury, Donghui Li, Alzheimer’s Disease Neuroimaging Initiative, Francesco Paolo Casale, Theofanis Karaletsos

**Affiliations:** Chan Zuckerberg Initiative, Redwood City, CA, USA; Institute of AI for Health, Helmholtz Zentrum München – German Research Center for Environmental Health, Neuherberg, Germany; Helmholtz Pioneer Campus, Helmholtz Zentrum München, Neuherberg, Germany

**Author notes:** Corresponding authors: Sayan Ghosal,; Theofanis Karaletsos. Part of the data used in preparation of this article were generated by the Alzheimer’s Disease Metabolomics Consortium (ADMC). As such, the investigators within the ADMC provided data but did not participate in the analysis or writing of this report. A complete listing of ADMC investigators can be found at: https://sites.duke.edu/adnimetab/team/.

## Abstract

Accurately predicting gene expression from DNA sequence remains a central challenge in human genetics. Current sequence-based models overlook natural genetic variation across individuals, while population-based models are restricted to variants observed within specific cohorts. Here, we present VariantFormer, a 1.2-billion-parameter transformer that predicts gene-level RNA abundance directly from personalized diploid genomes. Trained on 21,004 genome–transcriptome pairs from 2,330 donors, VariantFormer achieves state-of-the-art performance across both sequence- and population-based prediction tasks, while generalizing better to out-of-distribution contexts—including somatic mutation settings in cancer cell lines—and maintaining robustness across ancestries. Beyond expression prediction, VariantFormer improves eQTL effect size estimation compared to prior methods, with notable gains for lower-frequency and ancestry-specific variants. In applications to Alzheimer’s disease, VariantFormer gene embeddings prioritize likely causal genes and relevant tissue contexts, and in silico mutagenesis of known APOE alleles faithfully recovers known risk modifying effects. Together, these results establish VariantFormer as a scalable, diploid-aware framework for variant interpretation and personalized gene expression modeling across tissues and populations.

## 2 Introduction

Understanding how genetic variation modulates gene expression across tissues and individuals is fundamental to human genetics, disease biology, and precision medicine [21]. Gene expression is the most proximal yet measurable molecular consequence of genetic variation, yet predicting tissue-specific transcript abundance from an individual’s genome requires integrating multiple layers of biological complexity: the cis-regulatory grammar governing enhancer and promoter function [19, 26], long-range chromatin interactions [61], allele-specific dosage effects [53], and tissue-specific epigenetic programs [44, 18].

Two complementary paradigms have emerged to address this challenge: population level frequency based statistical genetics models [52] and DNA sequence based models [6]. Statistical genetics approaches like expression quantitative trait locus (eQTL) mapping [21] and Transcriptome-Wide Association Studies (TWAS) [22, 20] leverage population-scale genotype-expression correlations to quantify genetic effects on transcript abundance. By training gene-specific predictive models on common variants within cis-windows these methods capture individual-level expression variability and have proven highly successful for identifying disease genes through integration with genome-wide association study (GWAS) signals [52]. However, statistical genetics approaches are conceptually constrained by their reliance on training data: they cannot predict expression for genes absent from the training panel, variants not observed in training populations, or tissues lacking matched molecular phenotyping [36]. Each gene requires independent model training, precluding transfer of regulatory knowledge across loci and limiting their utility in understudied populations, rare diseases, and emerging single-cell contexts [58, 7].

In parallel, sequence-based models of genetic regulation have emerged to predict gene expression directly from DNA sequence. Early convolutional architectures (DeepSEA [60], Basset [28]) predicted chromatin accessibility and transcription factor binding from kilobase-scale windows, while subsequent models (Enformer [4], Borzoi [35], AlphaGenome [5]) scaled to megabase contexts (200 kb–1 Mb) for predicting epigenomic tracks and gene expression across hundreds of samples. Evolutionary-scale genomic foundation models (Evo [8], HyenaDNA [38], NT-Transformer [13], DNABERT [25]) further extended generalization by pretraining on billions of genomic tokens. However, these models share a critical limitation: they operate exclusively on reference genome sequences and lack native mechanisms to incorporate individual-specific diploid genotypes. While post-hoc approaches can score individual variants by comparing reference versus alternate allele predictions [54], this strategy does not account for LD structure, compound heterozygous configurations, or epistatic interactions between multiple cis-acting variants—features that fundamentally shape gene expression in diploid genomes [17, 23]. Importantly, the inability of reference-based models to accommodate personalized genomes limits their utility for equitable genomic medicine, as they cannot capture population-specific variation or rare variants enriched in underrepresented ancestries [34]. Recent hybrid approaches have explored fine-tuning foundation models on variant data [48], yet no existing framework fully integrates personalized diploid genomes, long-range cis-regulatory context, and tissuespecific conditioning for gene-level expression prediction at scale.

To address these challenges, we introduce VariantFormer, a 1.2-billion-parameter hierarchical transformer that predicts tissue-specific gene expression from personalized diploid genomes. VariantFormer integrates three key innovations: (1) it encodes individual genetic variants directly into DNA sequences using IUPAC ambiguity codes, enabling native modeling of heterozygous genotypes and haplotype effects; (2) it captures long range interactions with regulatory regions within megabase-scale cis-regulatory windows through mutation-aware transformer encoders; and (3) it employs hierarchical cross-attention to model regulatory influence of distal elements on gene bodies in a tissue-conditioned manner. We trained VariantFormer on 21,004 bulk RNA-seq samples from 2,330 donors across GTEx, 1000 Genomes, ENCODE, and ADNI cohorts, covering 54 tissues and 7 cell lines with paired whole-genome sequencing and expression data. This unified framework bridges statistical genetics and sequence-based modeling paradigms, enabling variant interpretation and disease risk assessment at scale.

We systematically evaluate VariantFormer across four axes: (1) *Gene expression prediction*: we benchmark performance against statistical-genetic baselines (e.g., genotype-based models) and sequence-based models (Enformer, Borzoi, Evo), measuring both cross-gene prediction within individuals and cross-individual prediction for each gene, stratified by gene type, tissue, and ancestry (Figures 2–3). (2) *Generalization*: we test out-of-distribution performance on high-mutation ENCODE cancer cell lines (Figure 2). (3) *Disease applications*: we assess Alzheimer’s disease risk stratification through tissue-specific MAGMA enrichment and supervised classification from learned gene embeddings (Figure 4). (4) *Variant-level validation and interpretability*: we validate effect predictions against independent eQTL catalogs (including low frequency variants and ancestry-stratified analyses), perform *in silico* APOE mutagenesis, and examine whether attention patterns align with chromatin accessibility (Figures 5-6).

## 3 Results

### 3.1 A hierarchical mutation-aware transformer for personalized gene expression prediction

To model genetic regulation of gene expression across genes, tissues, and individuals, we developed VariantFormer, a transformer trained on 21,004 paired whole-genome sequencing (WGS) and RNA-seq samples from 2,330 donors spanning four cohorts, 54 tissues, and seven cell lines (Figure 1). The 1.2-billion-parameter model was trained in two stages. In the first stage (Section 5.3), a mutation-aware module learned sequence representations trained to predict regulatory region activity across more than one million candidate cis-regulatory elements (cCREs) from ENCODE (Figure 7d). In the second stage (Section 5.4), these representations were integrated within a 25-layer hierarchical transformer that predicts gene-level expression from gene-centered transcription windows (≤ 300 kb downstream, 1 kb upstream; Figure 1a). Cross-attention between regulatory and gene modules enables explicit modeling of enhancer–promoter interactions across > 2 Mb of genomic context—substantially broader than prior frameworks. Finally, a tissue-conditioned prediction head outputs context-specific expression levels for each individual (Figure 1a). In this work we present two model variants: VariantFormer-PCG, trained on protein-coding genes, and VariantFormer-AG, trained on all annotated genes.

**Figure 1.**
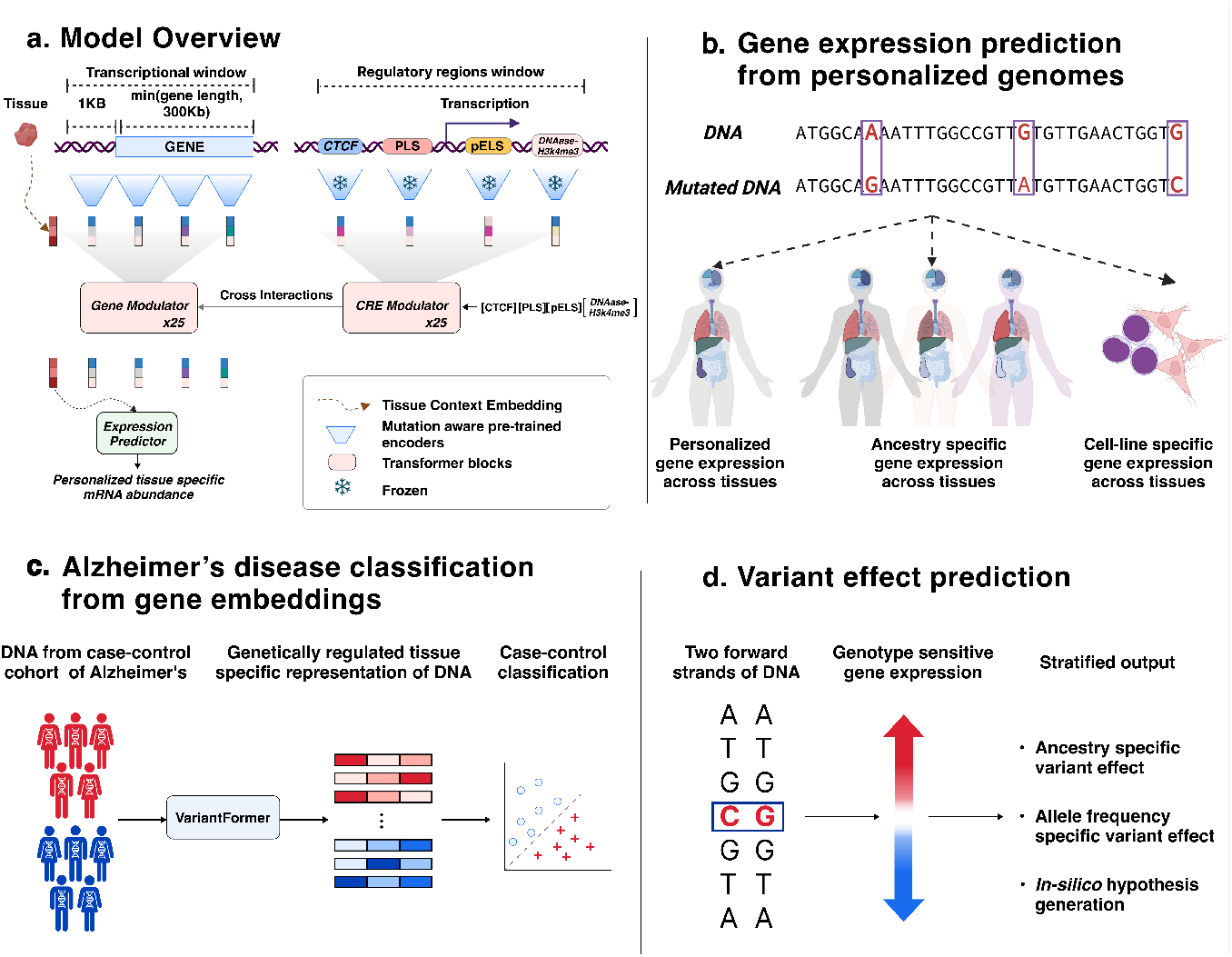
(a) Model architecture. VariantFormer integrates two complementary genomic windows for personalized gene expression prediction: (1) a cis-regulatory elements (CRE) window around the gene body, capturing distal regulatory regions; and (2) a gene transcription window, encompassing the gene body. The dual-transformer architecture comprises: (i) CRE modulators with pretrained frozen (snowflake symbol) encoders, and (ii) gene modulators with trainable encoders that attend to CRE representations via cross-attention. A tissue-specific context token (orange) conditions all layers, and its final representation feeds an expression predictor head (MLP) to output personalized, tissuespecific mRNA abundance. (b) Gene expression prediction from personalized genomes (highlighted in red boxes showing reference vs. mutated alleles). The model enables three complementary prediction modalities: (left) personalized gene expression prediction across diverse tissues; (middle) ancestry-specific gene expression prediction across populations; (right) gene expression prediction with somatic mutations, enabling out-of-distribution generalization to high-mutation contexts. (c) Alzheimer’s disease classification from gene embeddings. VariantFormer gene embeddings provide a genetically grounded representation for disease risk stratification. (d) Variant effect on gene expression and in silico mutagenesis. VariantFormer enables counterfactual analysis of genetic variants through genotype-sensitive expression prediction. This framework supports three complementary analyses: (i) ancestry-specific variant effects by embedding variants within population-matched haplotype backgrounds (red arrow, accounting for population-specific LD and regulatory contexts), (ii) allele frequency-stratified predictions enabling low frequency variant (MAF < 0.05) interpretation (blue arrow), and (iii) in silico hypothesis generation through computational genome editing, where disease alleles are reverted to reference states to isolate variant-attributable phenotypic effects while controlling for individual genetic backgrounds.

After training, we benchmarked VariantFormer across complementary tasks to assess predictive accuracy, generalization, and biological interpretability (Figure 1a-d. We evaluated performance in predicting gene expression across genes, tissues, and individuals; generalization to unseen genetic and somatic variation; and recovery of disease-relevant regulatory effects. Finally, we tested whether VariantFormer’s learned representations capture variant-level functional impacts and encode biologically coherent structure.

Our training and evaluation span four cohorts comprising 54 GTEx tissues, six ENCODE cell lines, one MAGE cell line, and an ADNI gene microarray cohort (details in Section 5.1; Section 5.6). Our held out data consists of 10% of the donors from each of the cohorts except for the ENCODE cell-lines. For ENCODE cell-lines we have 1 donor per cell-line so we removed Chromosome 19 from our training set and kept it as a held-out test set for evaluations. Figure 7b captures the data distribution and the train-test split of our datasets.

### 3.2 Performance on gene expression predictions tasks

We quantified the performance of the VariantFormer model on gene expression prediction task (Figure 2, 3) across all protein coding genes and non-coding genes on the held out test set (Section 5.6). Figure 2a shows the framework of the gene expression prediction task. We compared VariantFormer against the gold-standard genotype based RF model (genotype-RF) and three DNA based foundation models namely Borzoi, Enformer, and EVO 2. For the genotype based model we train a random forest model for each gene-tissue pair over the training data (Methods S1.1). On the other hand we trained tissue specific MLP heads [48] on top of the DNA embeddings of the foundation models (Section S1.2) for the gene expression prediction task. As shown in Figure 2b we evaluate the performance of each of the models across two complementary metrics, gene-correlations and subject correlations. Gene-correlation measures the ability of the model to capture variability across samples (donor-tissue pairs) for each gene, on the other hand subject correlation measures ability to capture the variability across genes.

**Figure 2.**
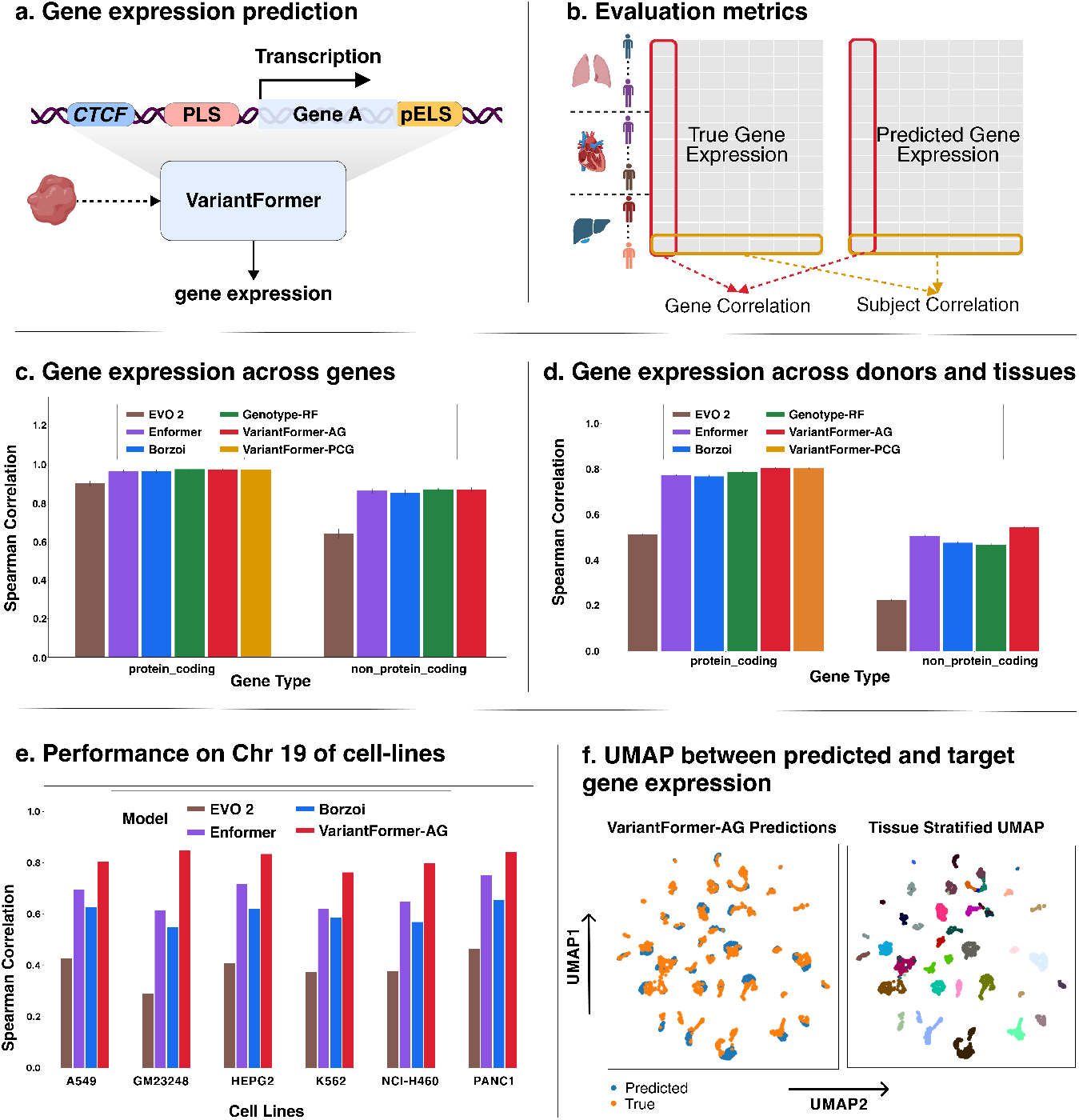
(a) Gene expression prediction task. VariantFormer takes personalized DNA sequences overlapping cCRE elements (spanning ±1 Mb) and transcription window containing genetic variants along with tissue context to predict gene-level expression (b) Evaluation framework. Gene correlation measures prediction accuracy across donors and tissues for individual genes (left, red); subject correlation measures prediction accuracy across all genes within individual samples (right, orange), capturing gene-to-gene expression variability. (c) Subject correlation performance across genes with 95% CI. Spearman correlation between predicted and observed expression across all genes for individual samples, stratified by protein-coding (n=18,439) and non-protein-coding (n=32,517) genes. (d) Gene correlation performance across donors and tissues stratified by gene type (proteincoding, n=18,439 genes; non-protein-coding, n=32,517 genes). (e) Performance on chromosome 19 in ENCODE cell lines. Spearman correlation between predicted and observed expression for all genes on chromosome 19 (held-out test set) across six ENCODE cell lines: five cancer cell lines (A549, HepG2, K562, NCI-H460, PANC1) and one lymphoblastoid cell line (GM23248). (f) UMAP visualization of predicted versus true gene expression. Left: UMAP projection of all samples colored by predicted (blue) versus true (orange) expression values. Right: Tissue-stratified UMAP colored by tissue type reveals that VariantFormer preserves tissue-specific expression patterns, with clear separation maintained across diverse tissue contexts.

**Figure 3.**
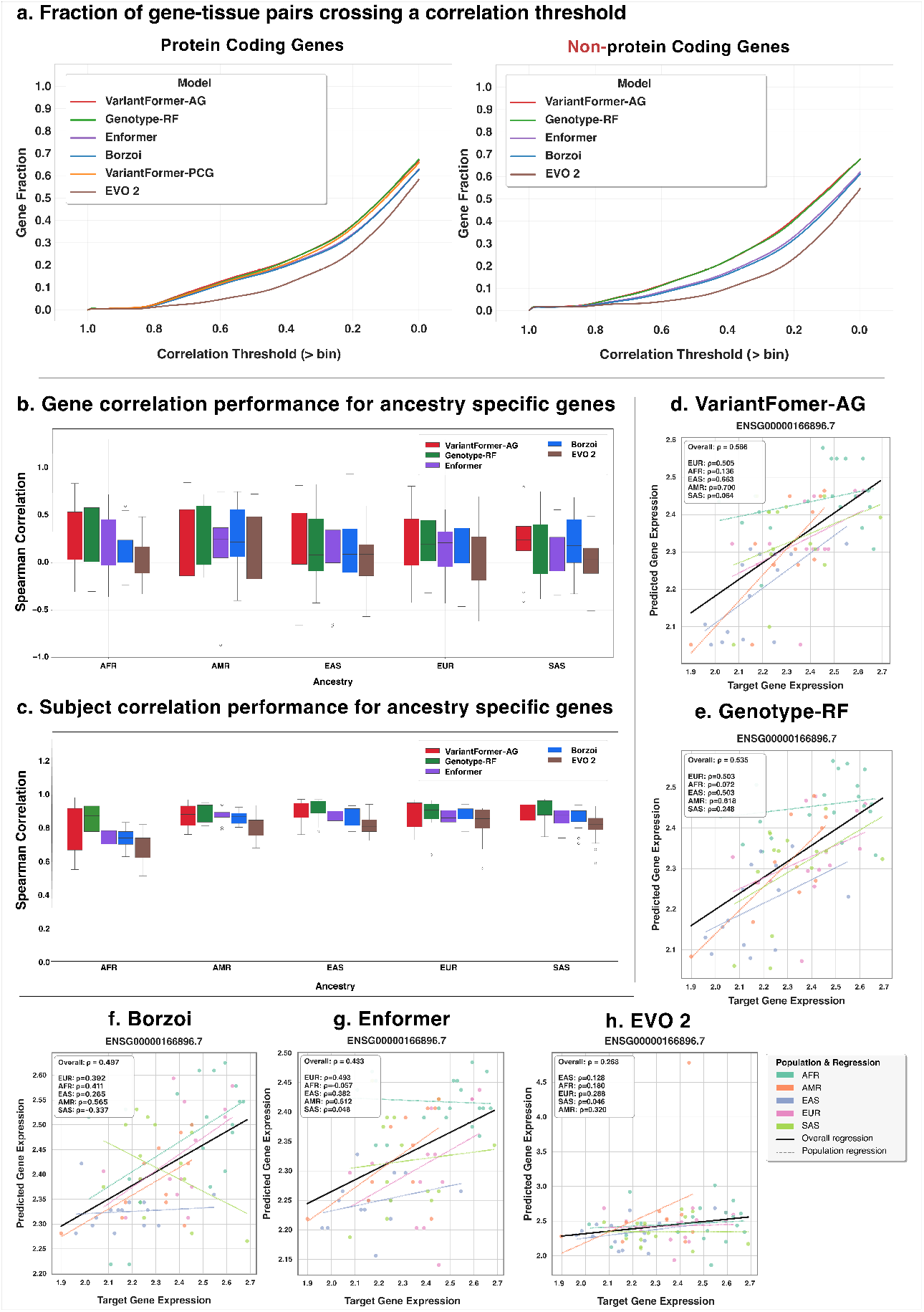
**(a)** Distribution of gene-tissue pair performance. The fraction of gene-tissue pairs exceeding correlation thresholds (x-axis) is shown for each model. **(b)** Gene correlation performance for ancestry-specific genes. Spearman correlation between predicted and observed expression across donors within each ancestry group for 23 ancestry-specific genes identified by one-way ANOVA (*p* < 10^−5^). Ancestries: African (*n* = 20), Admixed American (*n* = 11), East Asian (EAS, *n* = 14 donors), European (*n* = 14), South Asian (*n* = 14). **(c)** Subject correlation performance for ancestryspecific genes. Mean Spearman correlation across genes for individual samples within each ancestry group. **(d–h)** Predicted versus observed expression for ancestry-specific gene ENSG00000168896.7 across five genetic ancestries. Points are colored by ancestry, with dashed lines showing populationspecific regressions and solid black line showing overall regression across all populations.

**Figure 4.**
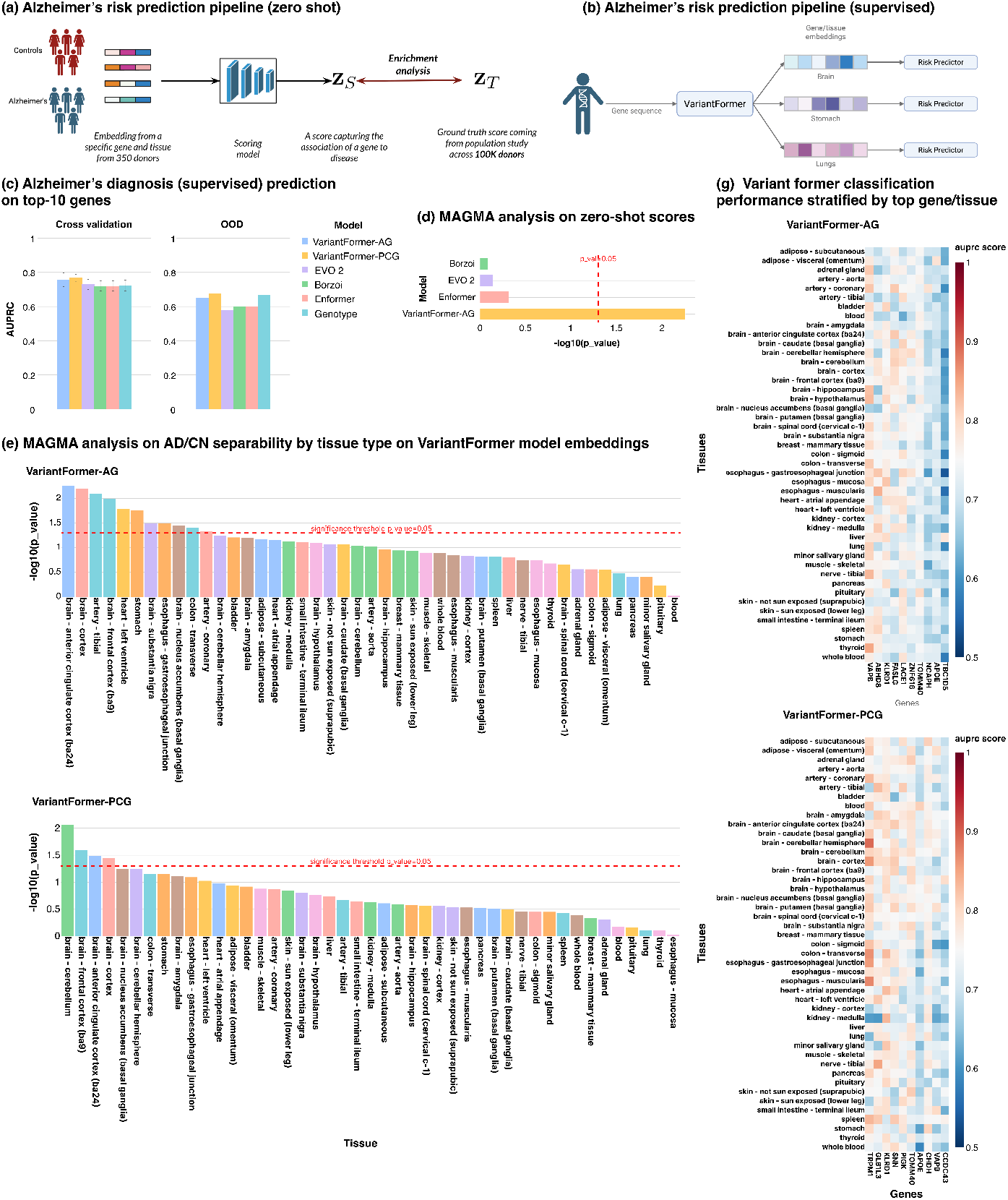
(a) Zero-shot MAGMA gene-property framework. Zeroshot scores are generated as distance measures. These scores are then tested with MAGMA enrichment analysis against GWAS risk statistics. (b) Supervised AD/CN classification with VariantFormer. (c) Comparison of VariantFormer (AG and PCG) with baseline models (EVO 2, Borzoi, Enformer, Genotype) on the supervised AD/CN classification task. AUPRC is reported for cross-validation (n = 330 donors) and out-of-distribution testing (n = 40 donors); VariantFormer-PCG performs best on the held-out test set. (d) MAGMA property significance for the top VariantFormer tissue (anterior cingulate cortex) versus baselines; the dashed line marks the significance threshold. (e) MAGMA tissue-property analysis of AD/CN separability using VariantFormer embeddings (AG, top; PCG, bottom). Multiple tissues exceed the threshold, with brain regions (e.g., cortex, cerebellum) showing the strongest enrichment. (g) Heatmaps of random-forest AUPRC across the top 500 gene–tissue pairs and corresponding top 10 genes for VariantFormer (AG and PCG). Higher values indicate stronger AD/CN discrimination; known AD loci—including APOE and its neighbor TOMM40—are prominent, highlighting tissues where the representations are most discriminative.

**Figure 5.**
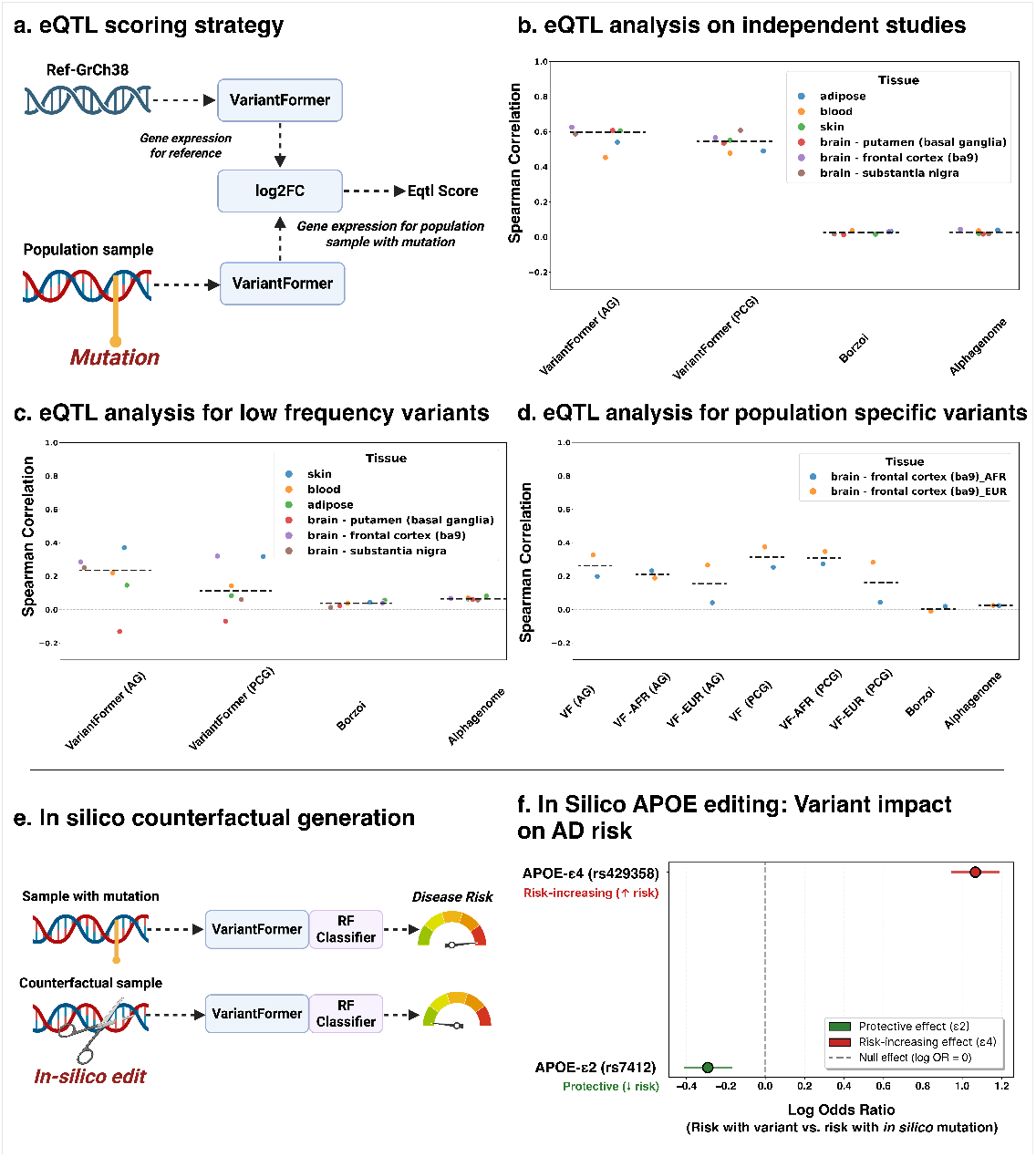
**(a)** eQTL scoring strategy. VariantFormer predicts gene expression from reference genome and from population-specific genomes containing the variant of interest. The log_2_ foldchange (log2FC) between variant-containing and reference predictions serves as the eQTL effect score. Gene expression predictions are aggregated across five ancestry-matched genomes (EUR, AFR, EAS, SAS, AMR) using an allele-frequency–weighted ensemble based on 1000 Genomes superpopulation frequencies. **(b)** eQTL validation across independent tissue-specific studies. Spearman correlation (*ρ*) between model-predicted variant effect scores and experimentally measured eQTL effect sizes from six independent tissues. Each point represents a tissue-specific eQTL dataset. **(c)** eQTL validation for low frequency variants (MAF < 0.05) based on spearman correlation **(d)** Population-specific eQTL validation. Spearman correlation for BrainSeq (brain frontal cortex) eQTLs stratified by ancestry-enriched variants: EUR-enriched (AF_EUR > 10%, AF_AFR < 5%) and AFR-enriched (AF_AFR > 10%, AF_EUR < 5%). For each variant, predictions were generated from ancestry-matched genomes (VF-EUR, VF-AFR) and compared to observed eQTL effects. **(e)** *In silico* counterfactual generation framework. APOE variant carriers from the ADNI cohort are analyzed under two genomic scenarios: (1) observed genotype with disease-associated variant, and (2) *in silico*-edited genome where the risk allele is computationally reverted to the neutral APOE-*ε*3 allele. VariantFormer generated tissue-specific gene embeddings are converted to AD risk scores using random forest classifiers (Section 6.2.2). **(f)** APOE variant impact on Alzheimer’s disease risk. Forest plot showing log odds ratios for APOE-*ε*4 (rs429358, risk-increasing) and APOE-*ε*2 (rs7412, protective) variants, comparing AD risk with observed variant allele versus *in silico ε*3-edited background. APOE-*ε*4 shows positive log odds ratio (log OR = +1.06 for VariantFormer AG, 95% CI: [0.95, 1.18]), indicating elevated AD risk consistent with established genetic architecture. APOE-*ε*2 shows negative log odds ratio (log OR = −0.29 for AG model, 95% CI: [−0.41, −0.17]), indicating protective effect against late-onset Alzheimer’s disease.

#### 3.2.1 Predicting gene expression across genes

Gene expression prediction across genes is the most widely used benchmark that measures the subject correlation for each sample and generates a population-level mean statistic. In Figure 2c, we showed the subject correlation performance of all models across protein-coding and non-protein-coding genes. Notably, both versions of VariantFormer achieve strong performance on this benchmark, with mean Spearman correlations of 0.97 for protein-coding genes and 0.87 for non-protein-coding genes (VariantFormer-AG), performing on par with or marginally better than the baseline models. The Genotype-Random Forest model achieved correlations of 0.97 (protein-coding) and 0.87 (non-protein-coding), while the DNA foundation models Enformer and Borzoi showed similar performance levels (0.96–0.97 for protein-coding and 0.86–0.87 for non-protein-coding genes). In comparison the EVO 2 finetuned on 8kb context, shows lower performance at 0.90 for protein-coding genes and 0.64 for non-protein-coding genes, which highlight that the pretraining paradigm significantly influence the downstream biological task.

The consistently high performance across all models, particularly for protein-coding genes where correlations exceed 0.96, indicates that this metric is highly saturated—an observation that aligns with recent findings in the field [23]. This saturation arises because measuring performance across genes primarily captures the large variability in baseline expression levels between different genes, which are relatively stable across individuals.

#### 3.2.2 Predicting population and tissue level variation in gene expression

Subject correlation across genes provides limited insight into the more challenging task of predicting individual-level and cross-tissue variation in gene expression. The subtle differences in expression levels across donors and tissues are substantially harder to predict, as they depend on complex interactions between genetic variants, regulatory context, and the individual’s genomic background. To assess this capability, we computed gene-level correlations that measure how well each model captures expression variability across different donors and tissue contexts for individual genes.

Figure 2d presents the gene correlation performance across protein-coding and non-protein-coding genes across all donors and tissues. In contrast to the saturated subject correlation metric, gene correlations reveal meaningful differences between models and demonstrate a more challenging prediction task. For protein-coding genes (n=18,439), VariantFormer achieves the highest performance with mean Spearman correlations of 0.804 (95% CI: 0.802-0.806) for VariantFormer-AG and 0.803 (95% CI: 0.801–0.805) for VariantFormer-PCG, substantially outperforming the baseline models. By comparison, the genotype based Random Forest (genotype-RF) achieved 0.787 (95% CI: 0.785–0.789), Enformer 0.774 (95% CI: 0.772–0.777), Borzoi 0.769 (95% CI: 0.767–0.771), and EVO 2 0.514 (95% CI: 0.512–0.517). This represents a relative improvement of 2.2% over genotype-RF and 3.9% over Enformer, demonstrating VariantFormer’s enhanced ability to capture individual-level variability in gene expression.

The performance gap becomes even more pronounced for non-protein-coding genes (*n* = 32,517). Here, VariantFormer-AG achieves a correlation of 0.544 (95% CI: 0.542–0.547), substantially outperforming Enformer at 0.507 (95% CI: 0.504–0.510), Borzoi at 0.476 (95% CI: 0.473–0.479), genotype-RF at 0.469 (95% CI: 0.466–0.472), and EVO 2 at 0.224 (95% CI: 0.221–0.227). This represents a 7.3% relative improvement over the second-best model (Enformer) and a 16.0% improvement over genotype-RF, highlighting VariantFormer’s superior capacity to model the complex regulatory landscape governing non-coding gene expression. EVO 2 finetuned on 8kb context window shows markedly lower performance (0.224) compared to other models underscores its limited capability in capturing the regulatory complexity of non-coding genes. Notably, the overall lower correlation values for non-protein-coding genes compared to proteincoding genes (0.22–0.54 vs 0.51–0.80) reflect the increased biological complexity and lower signal-to-noise ratio (Figure 7e) characteristic of these genes. Many non-coding RNAs exhibit tissue-specific [10], developmental stage-specific [50], or condition-specific [12] expression patterns that are challenging to capture from sequence alone. VariantFormer’s stronger performance in this regime suggests that its architecture effectively integrates sequence context, variant information, and tissue conditioning to better resolve these subtle regulatory signals.

Additionally, in Figure 2f we showed UMAP clustering plot of prediction and true gene expression counts. Qualitatively we see that VariantFormer faithfully captures the tissue specific separability and a high degree of overlap between the predicted and the true UMAP plots. Overall, these results establish that gene correlation is a more discriminative metric than subject correlation for evaluating gene expression prediction models, and demonstrate VariantFormer’s state-of-the-art performance in capturing individual-level transcriptional variation across diverse genetic backgrounds and tissue contexts.

#### 3.2.3 Predicting cross-donor variation in gene expression stratified by gene and tissue

In this analysis we focus on predicting cross-donor variation for each gene-tissue pair. This analysis addresses how accurately we can predict a specific gene’s expression that will vary across individuals with different genetic backgrounds within a specific tissue context. In this evaluation framework we stratify all the samples based on the tissues and compare the performance of the baseline model with VariantFormer. For comparison we combined the granular tissue like brain-putamen to brain such that for each gene we have enough samples to generate a robust estimate of the spearman rank correlations. To focus on genes where inter-individual variation is most biologically meaningful, we restricted our analysis to the top 3,000 highly variable genes (HVGs) within each tissue, ranked by expression variance across donors. These HVGs are enriched for genes whose expression is sensitive to genetic variation and environmental factors, making them more challenging to predict. For each gene-tissue pair, we computed the Spearman correlation between predicted and observed expression across all donors, yielding a distribution of per-gene correlation values that reflects model performance on individual genes rather than aggregate statistics

Figure 3a presents the cumulative distribution of gene-level performance across all models, showing the fraction of genes that exceed progressively stringent correlation thresholds. For protein-coding genes, VariantFormer and genotype-RF demonstrate uniformly better performance compared to the DNA-based models. At a correlation threshold of *ρ* > 0.4 VariantFormer-AG achieves 21.2% gene coverage, performing comparably to genotype-RF (20.9%) and better than Enformer (19.2%), Borzoi (18.8%), and EVO 2 (10.8%). In striking contrast, VariantFormer’s advantage becomes pronounced for non-protein-coding genes (Figure 3a), where the biological signal is substantially weaker and regulatory mechanisms more complex. At a relevant threshold of *ρ* > 0.5, VariantFormer-AG maintains predictive accuracy for 15.8% of non-coding genes, closely matching genotype-RF at 15.7% but substantially outperforming Enformer at 11.8%, Borzoi at 11.1%, and EVO 2 at 5.7%—representing a 34%, 42%, and 177% relative improvement over these DNA foundation models, respectively. This performance gap widens at lower thresholds: at *ρ* > 0.4, VariantFormer-AG predicts 21.5% of non-coding genes compared to only 16.4% for Enformer, 15.6% for Borzoi, and 9.2% for EVO 2, while at *ρ* > 0.2, VariantFormer-AG maintains 40.6% coverage versus 32.6% for Enformer, and 31.3% for Borzoi—representing 24%, and 30% relative improvements, respectively. On the other hand EVO 2 finetuned on a 8kb context window shows markedly lower performance across all thresholds highlights the challenges that large-scale DNA foundation models face without explicit variant encoding and tissue-specific conditioning.

The near-equivalence with genotype model for non-coding genes is notable, as genotype models are trained specifically on gene-tissue pairs with individualized genotype information. The results highlight VariantFormer’s ability to capture natural variations from DNA sequences alone, demonstrating its capacity to model variant interactions that transfers across diverse genetic backgrounds. The significant difference in performance patterns between protein-coding and non-coding genes reveals the inherent biological complexity of the regulatory landscape. Non-coding genes exhibit lower baseline correlations and wider performance gaps between models, reflecting their context-dependendencies. Many non-coding RNAs function in tissue-specific developmental processes [50] or respond to environmental stimuli [12], creating regulatory signals that are difficult to capture from sequence alone. VariantFormer’s superior performance in this challenging regime—particularly its substantial advantage over DNA foundation models—demonstrates its enhanced capacity to resolve complex regulatory interactions through integrated modeling of cis-regulatory elements, sequence variations and tissue contexts.

#### 3.2.4 Predicting gene expression of ancestry specific genes

Genetic ancestry significantly influences gene expression patterns through population-specific allele frequencies [49], linkage disequilibrium structures [56], and adaptive evolutionary pressures [43]. We evaluate VariantFormer’s performance across diverse genetic backgrounds, by assessing model performance on ancestry-specific genes. The ancestry specific gene were identified by performing one-way ANOVA across five major genetic ancestry groups in the heldout test MAGE dataset (Methods 6.1.3).

Figure 3c presents the distribution of subject correlations across all five ancestries for each model. Similar to Section 3.2.1 here we see saturation effects and all models demonstrate relatively strong and comparable performance. VariantFormer-AG achieves median subject-level correlations ranging from 0.796 (AFR) to 0.895 (EAS), with an overall mean of 0.858 across all ancestries. Genotype based Random Forest performs marginally better with a mean of 0.880, while Enformer and Borzoi achieve means of 0.826. However, EVO 2 shows substantially lower performance with subject-level correlations ranging from 0.557 (EUR) to 0.702 (SAS), with an overall mean of 0.627—representing a 37% relative decrease compared to VariantFormer. The relatively narrow performance range among VariantFormer, genotype-RF, Enformer, and Borzoi is mainly driven by large differences in mean expression between genes, which are stable across populations and relatively easy to predict, while EVO 2’s poor performance suggests fundamental limitations in capturing ancestry-specific expression patterns.

However, the more discriminative gene correlations reveal substantial and biologically meaningful differences in model performance. Figure 3b shows the distribution of gene-level correlations stratified by ancestry. VariantFormer-AG achieves the highest overall mean gene-level correlation of 0.244 (95% CI: 0.226–0.262), closely followed by genotype-RF at 0.237 (95% CI: 0.218–0.256). In comparison, Borzoi achieves 0.173 (95% CI: 0.157–0.189), Enformer 0.145 (95% CI: 0.128–0.162), and EVO 2 0.063 (95% CI: 0.043–0.083), representing 41%, 68%, and 287% relative improvements by VariantFormer over these DNA foundation models, respectively.

In Figure 3d-h we showed an example of the gene expression prediction for gene ENSG00000168896.7. As seen, VariantFormer achieves the best performance in terms of overall spearman correlation. In comparison the other DNA based models fail to capture the population specific trends which highlight their limitations to capture donor variability across populations.

The near-equivalence with genotype based Random Forest models is particularly notable, as these models are explicitly trained with population-specific genotype information for each gene-tissue pair and represent the current gold standard for incorporating genetic ancestry into expression prediction. However, a critical distinction emerges when examining population-level stability in gene-level correlations: VariantFormer demonstrates remarkably consistent median gene-level performance across ancestries (AFR: 0.329, AMR: 0.237, EAS: 0.199, EUR: 0.237, SAS: 0.239; overall median: 0.248, range: 0.199–0.329) compared to genotype-RF, which shows dramatically more variable gene-level performance (AFR: 0.409, AMR: 0.183, EAS: 0.081, EUR: 0.196, SAS: 0.152; overall median: 0.196, range: 0.081–0.409). Strikingly, genotype-RF gene-level correlation drops to 0.081 for EAS samples—representing an 80% decrease from its peak at AFR (0.409)—while VariantFormer maintains substantially tighter consistency across populations. This highlights that VariantFormer can extract ancestry specific information from sequence alone without requiring explicit ancestry labels or population-specific model training. The stability advantage is critical for clinical applications requiring reliable performance across diverse patient populations. These results establish that VariantFormer achieves state-of-the-art performance in predicting gene expression for ancestry-specific genes across diverse genetic backgrounds, substantially outperforming all DNA foundation models while maintaining superior generalizability and cross-population stability compared to population-specific genotype-RF models. This capability is essential for equitable genomic medicine, as it enables accurate and consistent prediction of gene expression and variant effects across all populations, including understudied groups where population-specific training data may be limited or unavailable.

#### 3.2.5 Gene expression prediction across gene in presence of somatic mutations

Somatic mutations present a unique challenge for gene expression prediction models due to their out-of-distribution characteristics. To evaluate model performance in this challenging regime, we leveraged ENCODE data consisting of six cell lines—five cancer cell lines (K562, A549, HepG2, Panc1, NCI-H460) and one lymphoblastoid cell line (GM23248)—with paired whole-genome sequencing (WGS) and RNA-seq data. Somatic variants were identified and distinguished from germline variants using DeepVariant and DeepSomatic (Section S1.4). The somatic variant burden in these cell lines ranges from 0.1:1 to 1.2:1 relative to germline variants, depending on the cell line and cancer type (Figure 7i). Our held-out evaluation set consists of all genes on chromosome 19 that corresponds to the cell-lines. These genes were removed from the training data and used to assess model generalization in high-mutation contexts. This experimental design evaluates a model’s ability to handle somatic variants not present in the training distribution.

This framework also exposes a fundamental limitation of traditional genotype-based approaches. Genotype-based models rely on pre-computed weights derived from specific variant sets observed during training. Because these models cannot dynamically incorporate novel variant combinations or generalize to unseen genes, they are architecturally incapable of making predictions in this setting and are therefore excluded from this analysis. This represents a critical practical constraint: genotype based models cannot be applied to contexts with extensive somatic variation, low frequency variants, or any scenario requiring prediction for genes or variant combinations outside the training distribution.

Figure 2e presents the Spearman correlation between predicted and observed gene expression across all chromosome 19 genes for each cell line. The performance advantage is consistent across diverse cell line types. VariantFormer achieves the highest correlations in GM23248 (0.848) and Panc1 (0.84), followed by HepG2 (0.834), A549 (0.805), NCI-H460 (0.800), and K562 (0.763). In comparison, Enformer correlations range from 0.613–0.752, while Borzoi ranges from 0.549–0.655. The performance gap highlights that VariantFormer can explicitly model individual-specific variants through IUPAC encoding integrated directly into the sequence representation, allowing it to learn how specific variant combinations alter regulatory logic.

### 3.3 Tissue-specific gene embeddings enable Alzheimer’s disease risk stratification

In this task we evaluate VariantFormer’s ability to capture Alzheimer’s disease (AD) disease-relevant biological signals in the learned latent representation space from the tissue-conditioned personal genomes. We analyzed 370 donors from the Alzheimer’s Disease Neuroimaging Initiative (ADNI) [41, 55] comprising 215 AD cases and 155 cognitively normal (CN) controls. We performed two complementary evaluations: **(1)** zero-shot MAGMA enrichment analysis to test whether embedding-derived gene scores correlate with independent GWAS signals without training classifiers, and **(2)** supervised classification to quantify predictive accuracy of gene and gene-tissue embeddings for AD risk.

#### 3.3.1 Evaluation of VariantFormer zeroshot scores using MAGMA enrichment analysis

To validate that VariantFormer’s learned embeddings capture genetically grounded disease signals without supervised training, we performed zero-shot gene-property enrichment analysis using MAGMA [16]. Critically, this evaluation requires no classifier training—we derive per-gene AD/CN separability scores directly from the embedding geometry across donors (Figure 4a; Section 6.2.1), then test whether these scores correlate with independent GWAS gene-level association statistics. This provides an orthogonal validation of biological coherence, less susceptible to overfitting than supervised methods.

VariantFormer’s unique tissue-conditioning allows us to identify tissue specific susceptibility of genetic risk. In contrast, baseline DNA models (Borzoi, Enformer, EVO 2) produce tissue-agnostic representations. Figure 4d compares model performance using anterior cingulate cortex—VariantFormer’s top-performing tissue—as a representative example. Other than VariantFormer, *no baseline model exceeds the genome-wide significance threshold* (*p* < 0.05, dashed line).

VariantFormer-AG and VariantFormer-PCG identify multiple tissues crossing the significance threshold, with strongest enrichment in brain structures including anterior cingulate cortex, cerebellar hemisphere, cortex, and frontal cortex—regions with well-established roles in AD pathology [57, 24, 27]. Notably, several peripheral tissues (stomach, heart, colon) also show enrichment, potentially reflecting systemic inflammatory or vascular components of AD pathogenesis [11, 46]. The tissue-specific enrichment pattern demonstrates that VariantFormer’s learned representations capture not only which genes contribute to disease risk, but the tissue contexts in which this genetic risk manifests.

#### 3.3.2 Supervised classification

In this task we leverage embeddings from four baseline models—EVO 2, Borzoi, Enformer, and Genotype—and compare them to both VariantFormer variants (AG and PCG). A pipeline sketch is shown in Figure 4b. In the Genotype model, each gene is represented by a genotype matrix analogous to the previous genotype-based model used for transcription prediction. For all baseline models, we train a random forest (RF) classifier on per-gene representations (Section 6.2.2); for VariantFormer, which explicitly conditions on tissue, we train separate classifiers for each gene–tissue pair. Data splits are performed at the donor level (train/eval/test). RF hyperparameters are selected via cross-validated on the train/eval partition. Genes (or gene–tissue pairs) are then ranked by AUPRC score, and test-set metrics are reported for the top K (K = 10) score, reflecting the biological expectation that only a subset of genes contribute strongly to AD signal. The same procedure is applied consistently across all baseline models. Figure 4c summarizes model performance in terms of AUPRC for both cross-validation (330 donors) and out-of-distribution testing (40 donors), where the VariantFormer-PCG model outperforms all baselines on the held-out cohort. Because we train an RF classifier for each gene–tissue pair, we can quantify classification performance at fine granularity, identifying where AD/CN discrimination is most pronounced. Figure 4g presents a heatmap of AUPRC values across the top 10 genes, revealing concentrated signal at known AD-associated loci—such as APOE [47] and TOMM40 [45]—and highlighting tissues where the learned representations are most discriminative.

### 3.4 Investigating in silico mutation and eQTL analysis using VariantFormer

In this task we assess whether VariantFormer captures biologically meaningful regulatory effects of genetic variants. We employed two complementary validation strategies: (i) benchmarking predicted expression changes against independent eQTL effect sizes across six tissue-specific datasets spanning diverse allele frequencies and ancestries, and (ii) performing in silico mutagenesis on APOE variants to test whether counterfactual predictions recapitulate known genotype-disease associations in Alzheimer’s disease.

For eQTL analysis, we estimated variant impact by contrasting VariantFormer predictions under two scenarios: a reference genome baseline and population-specific genomes (EUR, AFR, EAS, SAS, AMR) with the focal allele introduced. This approach embeds variants within realistic ancestry-matched haplotype and linkage disequilibrium contexts, enabling the model to capture population-specific *cis*-regulatory interactions. We converted expression predictions into log_2_ fold-change scores and aggregated population-specific predictions using an allele-frequency–weighted ensemble (Section 6.3.1).

For *in silico* mutagenesis, we applied VariantFormer to patient genomes from the Alzheimer’s Disease Neuroimaging Initiative (ADNI) cohort (Section 5.1.9), generating predictions under observed genotypes and computationally edited genomes where APOE risk alleles (*ε*2, *ε*4) were reverted to the neutral *ε*3 allele. This counterfactual design isolated variant-attributable effects while controlling for individual genetic backgrounds. We converted resulting gene embeddings into tissue-specific AD risk scores using random forest classifiers trained on the Alzheimer’s classification task (Section 6.3.2). Together, these analyses evaluate VariantFormer’s capacity to quantify variant regulatory impact with the potential to informing precision medicine applications.

#### 3.4.1 eQTL analysis in three independent studies

We benchmarked predicted expression changes against independent eQTL effect sizes from the eQTL Catalogue [29]. We compared Variantformer against SOTA model Borzoi and AlphaGenome (see Section S1.3). We selected six tissue-specific datasets (Section 5.1.7) spanning peripheral tissues from TwinsUK (adipose, blood, skin) and brain regions from Braineac2/BrainSeq (substantia nigra, frontal cortex BA9, putamen). Within each study we selected statistically significant variant–gene pairs (study-specific thresholds; see Section 5.1.7) and restricted to variants that fall within the context windows of VariantFormer, AlphaGenome, and Borzoi (variants are provided in Supplementary Table 1). We then computed Spearman correlation (*ρ*) between reported eQTL slopes and model-derived scores.

VariantFormer consistently outperformed DNA-only baseline models (Borzoi and AlphaGenome) across all six independent datasets (Figure 5)b. The allele-frequency–weighted ensemble score—which aggregates predictions across five ancestry-matched genomes (EUR, AFR, EAS, SAS, AMR) weighted by 1000 Genomes super-population frequencies—achieved the highest median correlation with eQTL effect sizes (*ρ* = 0.6), compared to Borzoi (*ρ* ≈ 0) and AlphaGenome (*ρ* ≈ 0). This advantage held across both peripheral and brain tissues.

Importantly, these eQTL datasets were entirely independent: discovery cohorts, expression quantification methods (RNA-seq processing pipelines), and population structures all differed from those used during model training. The consistent replication demonstrates that VariantFormer’s integration of personalized genomic context and long-range cis-regulatory interactions generalizes beyond training data to capture true regulatory mechanisms, even when discovery conditions vary substantially.

#### 3.4.2 eQTL analysis for Low-frequency (MAF-stratified) variants

Most disease-associated variants are of low to moderate frequency, yet statistical eQTL discovery power drops sharply below 5% minor allele frequency (MAF). To assess whether VariantFormer retains predictive accuracy in this challenging regime, we repeated the eQTL replication analysis after restricting to low-frequency variants (MAF < 0.05; Section 5.1.7).

VariantFormer maintained robust correlations with eQTL effect sizes for low frequency variants across both TwinsUK and brain tissues (median *ρ* = 0.20), substantially outperforming Borzoi (median *ρ* = 0.04) and AlphaGenome (median *ρ* = 0.06; Figure 5c). The wide performance gap across all six tissue datasets shows the generalization capabilities of VariantFormer that is not tissue or cohort-specific.

These results have important implications for precision medicine: the majority of non-coding GWAS hits and putatively causal fine-mapped variants fall in the low-to-moderate frequency range where direct experimental characterization is costly and eQTL power is limited. VariantFormer’s ability to prioritize functional candidates among low frequency regulatory variants—including those below standard eQTL detection thresholds—provides a computational avenue for accelerating follow-up experiments and interpreting individual genomes.

#### 3.4.3 eQTL analysis for ancestry specific variants

To test whether VariantFormer’s ancestry-matched predictions capture population-specific regulatory effects, we analyzed brain eQTL data from BrainSeq (total donors = 479, EUR donors = 231, AFR donors = 195, SAS donors = 1, Unassigned = 52) after stratifying variants by allele frequency in 1000 Genomes: EUR-enriched (AF-EUR > 10%, AF-AFR < 5%) and AFR-enriched (AF-AFR > 10%, AF-EUR < 5%) variants. For each variant, we compared VariantFormer predictions derived from EUR versus AFR genomes (VF-EUR, VF-AFR) against observed eQTL effect sizes.

VariantFormer’s ancestry-matched predictions demonstrated population-specific concordance with eQTL signals (Figure 5d). For AFR-enriched variants, the AFR-genome based prediction substantially outperformed the EUR-genome based prediction (VF-AFR: *ρ* = 0.27 vs. VF-EUR: *ρ* = 0.04 for PCG model;

*ρ* = 0.23 vs. 0.04 for AG model), a 6–7 fold improvement. For EUR-enriched variants, VF-EUR achieved *ρ* = 0.27 (AG) and 0.28 (PCG) compared to VF-AFR at *ρ* = 0.19 (AG) and 0.35 (PCG), demonstrating variable ancestry sensitivity across model architectures. The sensitivity to the model version stems from the fact that VariantFormer (AG) is trained over a larger set of genes over a significantly longer training period (Section 5.5) making it a better model for the eQTL task. Finally, we note that the allele-frequency–weighted ensemble consistently achieved strong performance across both variant sets (EUR: *ρ* = 0.33 AG, 0.38 PCG; AFR: *ρ* = 0.20 AG, 0.25 PCG), dynamically integrating population-specific information. In contrast, reference-genome models showed no ancestry discrimination: Borzoi achieved *ρ* = −0.01 (EUR) and 0.02 (AFR), while AlphaGenome showed uniform low correlation (*ρ* ≈ 0.025 for both).

These results demonstrate that personalized genome modeling enables ancestry-aware variant interpretation, with the allele-frequency–weighted ensemble providing robust predictions across populations—a critical advantage for genomic medicine in globally diverse cohorts.

#### 3.4.4 In silico mutation of APOE allele

In an exploratory study we evaluate VariantFormer’s ability to generate counterfactual predictions (see Section 6.3.2) by *in silico* editing the genome. We performed the counterfactual analysis on APOE variants— the strongest genetic risk factors for Alzheimer’s disease. Using AD patients from our held-out test cohort (*N* = 12 APOE-*ε*4 carriers, *N* = 1 APOE-*ε*2 carriers), we generated personalized predictions under two genomic scenarios: (i) the patient’s observed genotype and (ii) an *in silico*-edited genome in which the APOE risk allele was reverted to the neutral *ε*3 allele (rs429358-C→T for *ε*4; rs7412-T→C for *ε*2). This approach isolated the regulatory contribution of each APOE variant while holding constant the individual’s genome-wide genetic background.

For each genotype configuration, VariantFormer generated the APOE gene embeddings across 13 brain tissues. We converted tissue-specific gene embeddings into AD risk scores using random forest classifiers trained on Alzheimer’s disease classification task (Section 6.2.2). To quantify the magnitude and direction of variant effects, we computed log odds ratios comparing the risk with the observed variant allele versus the *in silico ε*3-edited background.

Figure 5e presents a forest plot showing log odds ratios for both APOE variants across both model versions. The results demonstrate that VariantFormer’s predictions align perfectly with the established genetic architecture of APOE alleles:

##### APOE-*ε*4 increases predicted AD risk

The *ε*4 variant (rs429358-C) shows a positive log odds ratio, indicating elevated AD risk relative to the *ε*3 background (log OR = +1.06 for VariantFormer-AG, 95% CI: [0.95, 1.18]; Figure 5e). This recapitulates the well-established population-level association between *ε*4 and increased AD susceptibility, with consistent directionality across all brain tissues [47].

##### APOE-*ε*2 decreases predicted AD risk

Conversely, the *ε*2 variant (rs7412-T) shows a negative log odds ratio, indicating reduced AD risk compared to *ε*3 (log OR = −0.29 for AG model, 95% CI: [−0.41, −0.17]; Figure 5e). This is consistent with *ε*2’s known protective effect against late-onset Alzheimer’s disease [47]. The symmetric response to risk-increasing (*ε*4) and protective (*ε*2) alleles demonstrates that VariantFormer’s predictions align with established genetic architecture rather than reflecting model bias. These results validate that VariantFormer integrates genomic variation, tissue-specific regulatory context, and disease-relevant expression patterns to produce biologically coherent risk predictions. The ability to perform *in silico* mutagenesis—replacing disease alleles with reference alleles while preserving an individual’s genetic background—provides a computational framework for dissecting variant contributions to complex disease phenotypes without requiring matched experimental perturbations.

### 3.5 Generalization capabilities through learned attention and embeddings

Transformer-based sequence models derive their predictive power from two core architectural components: attention modules, which dynamically weight input features based on context, and embeddings, which encode sequence information into dense vector representations. To evaluate whether VariantFormer has learned generalizable biological principles rather than task-specific correlations, we assessed both components on orthogonal validation tasks for which the model received no explicit training. Specifically, we tested whether (i) attention weights recapitulate experimentally measured chromatin accessibility patterns, indicating that the model has learned the regulatory grammar governing tissue-specific gene expression, and (ii) gene embeddings capture tissue context and function, demonstrating that learned gene representations encode fundamental properties of the gene.

#### 3.5.1 Attention patterns recover chromatin accessibility landscapes

We first evaluated whether VariantFormer’s attention mechanism identifies biologically functional regulatory regions by comparing attention weights against DNase-seq measurements of chromatin accessibility. DNase hypersensitivity marks genomic loci where chromatin is open and accessible to transcriptional machinery, providing an independent biochemical readout of regulatory activity. If VariantFormer has learned meaningful regulatory grammar, regions receiving high attention should correspond to accessible chromatin regions.

We analyzed a random subset of 50 protein-coding genes in adrenal gland tissue (UBERON:0002369). DNase-seq chromatin accessibility data for adrenal gland tissue were obtained from ENCODE (accession ENCFF632QUC) in bigWig format. Gene annotations including gene start sites and strand orientation were extracted from GENCODE v24 basic annotations aligned to the GRCh38 reference genome. For each gene, we passed the reference genome sequence through VariantFormer conditioned on adrenal gland tissue context, extracting cross-attention weights from the gene modulator layers attending to *cis*-regulatory element (CRE) embeddings. We then computed Spearman correlations between attention scores across cCREs and corresponding DNase-seq signal intensities, quantifying the concordance between model attention and experimental accessibility (Figure 6a).

**Figure 6.**
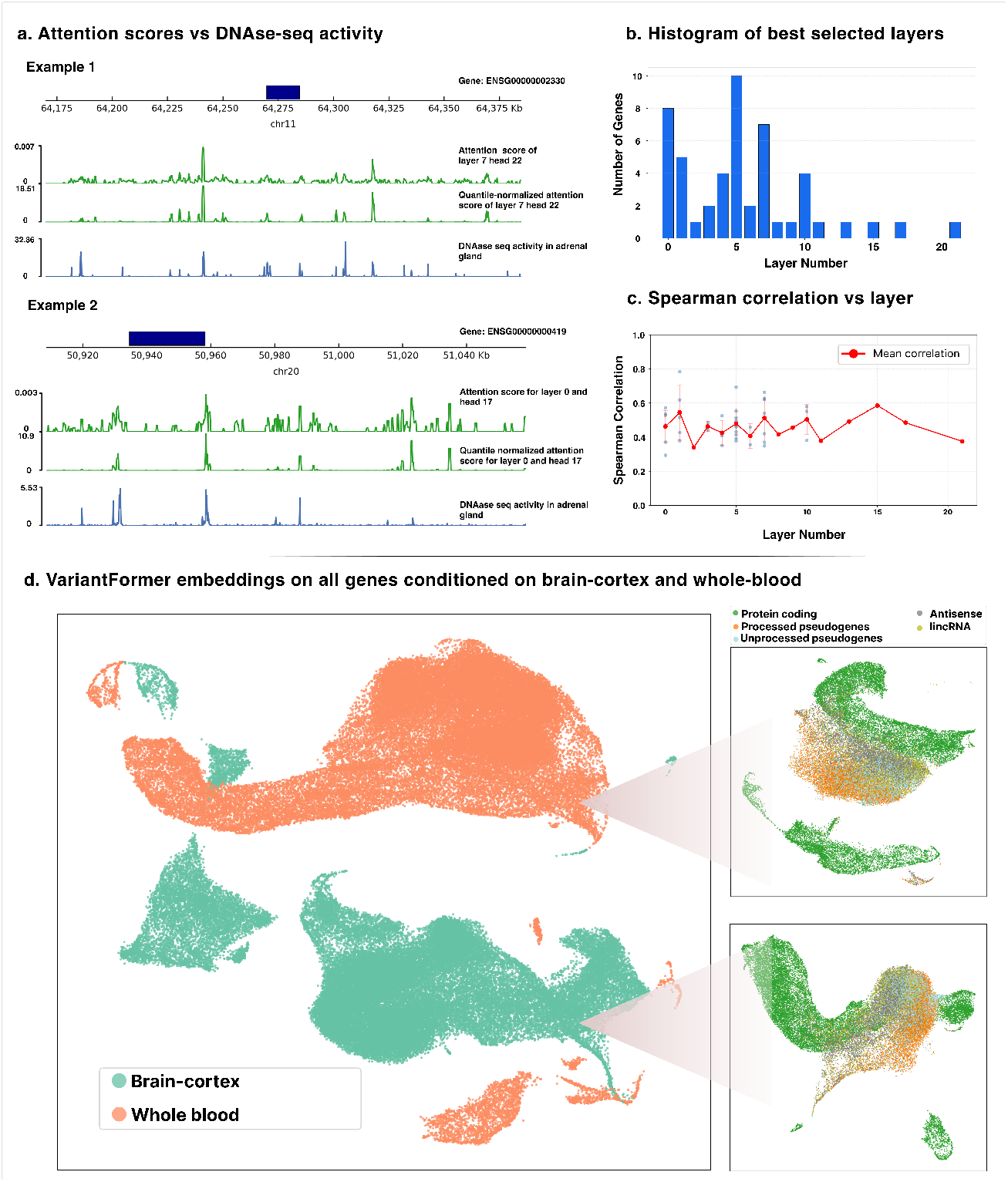
**(a)** Attention scores recapitulate chromatin accessibility landscapes. Two representative examples showing alignment between VariantFormer cross-attention weights and DNase-seq chromatin accessibility in adrenal gland tissue. In each gene panel top track shows the attention scores coming from the optimal head and layer, middle track shows the DNAse-seq signal. Gene bodies are shown as dark blue bars above each panel. **(b)** Distribution of optimal transformer layers across genes. Histogram showing the frequency with which each transformer layer (0–24) is picked as the optimal layer for 50 randomly selected protein-coding genes in adrenal gland tissue. Layer 5 emerges as the most frequently optimal layer (10 of 50 genes, 20%). **(c)** Optimal layer-specific correlation with chromatin accessibility. Mean Spearman correlation (±95% CI) between attention scores and DNase-seq signals across all 50 genes as a function of transformer layer depth. **(d)** Gene embeddings capture tissue-level organization. UMAP projection of gene embeddings for two tissues: brain-cortex and whole blood. Colors indicate tissue of origin. Higher resolution within each tissue highlights clustering of protein coding genes, pseudo-genes, and lincRNAs.

VariantFormer attention patterns showed substantial concordance with DNase-seq signals, with optimal layer-head combinations (Methods 6.4.1) achieving mean Spearman correlation of *ρ* = 0.48 (range: 0.29–0.78) across 50 genes (Figure 6c). This demonstrates that VariantFormer learns to attend to regulatory regions that are experimentally accessible. Notably, performance varied systematically across transformer layers, revealing hierarchical processing of regulatory information (Figure 6b,c). Early layers (0–10) showed strong to moderate correlations, with layer 5 emerging as the most frequently optimal layer across genes (10 of 50 genes, 20%; mean *ρ* = 0.479). The layer-specific correlation distribution revealed a characteristic pattern that VariantFormer’s architecture hierarchically refines regulatory representations. Lower layers capture local chromatin features and individual element activity, while deeper layers progressively integrate these signals into gene-level expression predictions that abstract away from specific accessibility patterns. Gene-specific analysis showed that optimal attention layers vary systematically (Figure 6b), suggesting that different genes employ distinct regulatory architectures—some relying on proximal elements resolved in early layers, others requiring distal enhancer integration captured in mid-tier layers.

Critically, this concordance was observed using only the reference genome sequence without individual-specific variants, demonstrating that VariantFormer’s base representations capture fundamental principles of tissue-specific gene regulation. While the model was trained to predict gene expression the attention patterns emergently align with DNase-seq signals through self-supervised learning. This demonstrates that VariantFormer’s attention mechanism takes advantage of the regulatory grammar of gene expression— learning which genomic contexts are functionally relevant for transcriptional control.

#### 3.5.2 Gene embeddings span biologically structured latent space

To evaluate whether VariantFormer’s learned representations capture fundamental biological organization, we extracted gene embeddings for all genes conditioned on brain cortex and whole blood, projecting them into two-dimensional space using UMAP (Fig. 6d). The gene embedding are obtained directly from the reference genome (Section 3.5.2). Gene embeddings from the two tissues form completely separate clusters, confirming the model has learned tissue-specific regulatory programs. Within each tissue, we colored genes by gene type (GENCODE biotypes). Most strikingly, gene type annotations reveal separation of protein coding genes, psudogenes and lincRNAs. Together with the attention concordance analysis, these results establish that VariantFormer’s learned representations encode interpretable biological principles of gene regulation beyond task-specific expression prediction.

## 4 Discussion

VariantFormer provides a unified approach to predict gene expression variation across genes, tissues, and individuals from personalized diploid genomes. VariantFormer bridges statistical genetics and deep learning paradigms by integrating diploid sequences, mutation-aware transformers, and learnable tissuespecific registry tokens. Critically, our pretraining strategy—grounding sequence encoders in tissue-specific chromatin accessibility prediction using donor-specific ENTEx data—enables the model to learn regulatory grammar relevant to downstream gene expression tasks. The asymmetric training design, where CRE encoders remain frozen while gene encoders adapt, mirrors biological hierarchy: distal regulatory elements establish stable cell-type identity through chromatin states, while gene-proximal regions integrate these signals with transcriptional machinery.

VariantFormer demonstrates state-of-the-art performance on discriminative metrics capturing individual-level variability, with substantial advantages for non-coding genes (correlation 0.544 versus 0.507 for Enformer, representing 7.3% improvement) where regulatory mechanisms are most complex. Critically, it generalizes to out-of-distribution contexts where existing approaches architecturally fail. On ENCODE cell lines with high somatic mutation burden, VariantFormer maintains correlations of 0.76–0.85 versus 0.55–0.75 for reference-based models; genotype based models cannot make predictions in this regime as they rely on precomputed variant-expression weights that do not transfer to novel mutation patterns. For ancestry-specific genes, VariantFormer exhibits superior cross-population stability (performance range 0.199–0.329 across five ancestries) compared to genotype-RF (0.081–0.409, representing 80% variation from peak to trough), with particularly striking differences in East Asian samples where genotype-RF correlation drops to 0.081 while VariantFormer maintains 0.199. This demonstrates that sequence-based learning with variant encoding captures population-specific regulatory variation without requiring ancestry-specific retraining—a critical capability for equitable genomic medicine where models trained predominantly one population and show degraded performance in underrepresented populations [30].

Beyond predictive performance, we demonstrate for the first time that learned gene embeddings from sequence-based models capture disease-relevant biological signals without explicit disease-specific training. Tissue-conditioned embeddings enable Alzheimer’s disease risk stratification through MAGMA enrichment, where gene-level separability scores derived purely from embedding geometry correlate with independent GWAS statistics, with strongest enrichment in cortical brain regions established in AD pathology [57, 24]. Supervised classification from embeddings outperforms genotype-based baselines, with APOE and TOMM40 emerging as top discriminative loci. Critically, *in silico* APOE mutagenesis—computationally reverting risk alleles to neutral backgrounds while preserving individual genetic context—correctly recapitulates established disease architecture: *ε*4 increases predicted risk (log OR = +1.06, 95% CI: [0.95, 1.18]) while *ε*2 shows protective effects (log OR = −0.29, 95% CI: [−0.41, −0.17]). This symmetric response demonstrates mechanistic faithfulness rather than model bias. Variant-level validation against independent eQTL catalogs demonstrates strong replication (median *ρ* = 0.6) extending to low-frequency variants (MAF < 0.05: *ρ* = 0.20 versus ∼0 for reference-based models) and, for the first time in any sequence-based model, ancestry-specific concordance: AFR-enriched variants show 6–7 fold higher correlation when predicted from AFR-matched genomes versus EUR genomes (0.27 versus 0.04), while reference-based models show no ancestry discrimination. These ancestry-aware predictions address a fundamental equity challenge in variant interpretation, enabling accurate functional assessment across diverse populations including those underrepresented in eQTL discovery studies.

Mechanistic interpretability analyses reveal that VariantFormer learns generalizable regulatory principles. Cross-attention weights show substantial concordance with DNase-seq chromatin accessibility (mean *ρ* = 0.48 across 50 genes), with hierarchical layer structure where early layers capture local chromatin features. This emergent alignment with biochemical measurements demonstrates that the model internalizes which genomic contexts are functionally relevant for transcriptional control. Additionally the gene embeddings encode fundamental gene properties: when used as a gene level encoding vector they capture tissue and gene-type specific clustering.

Despite its advances, VariantFormer shares multiple challenges. First, VariantFormer’s context window is restricted to cis-regulatory elements and transcription windows within megabase-scale neighborhoods, precluding direct modeling of trans-acting factors, cross-gene regulatory networks, or chromosomal interactions beyond sequence proximity. Second, training was limited to autosomal chromosomes, excluding sex chromosomes and their dosage compensation mechanisms. Third, while the model captures regulatory effects mediated through shared cis-elements between genes, it cannot explicitly model direct gene-gene interactions or transcriptional cascades. Future work could address these limitations by integrating Hi-C-derived chromatin contact maps to capture three-dimensional genome organization, extending to sex chromosomes with appropriate normalization strategies, and incorporating trans-regulatory signals through multi-gene context windows or graph neural network architectures that explicitly model regulatory cascades. The model predicts steady-state expression but not dynamic responses to environmental perturbations or developmental transitions; incorporating temporal context could extend to longitudinal studies. Additional directions include scaling to single-cell resolution through fine-tuning on cell-type atlases. By enabling accurate, tissue-specific prediction of gene expression from personalized genomes—including low frequency variants and diverse ancestries—VariantFormer provides a computational foundation for variant interpretation, disease risk assessment, and biomarker prioritization.

## 5 Methods

### 5.1 Data acquisition and processing

#### 5.1.1 Reference genome and genomic coordinates

VariantFormer is trained with WGS, RNA-seq and cis-regulatory elements data, all mapped to human genome assembly GRCh38 (GRCh38_no_alt_analysis_set_GCA_000001405.15)[21]. Additionally, we restricted analysis to autosomes only (Chr1-Chr22). For genomic coordinates and gene annotation, we used GENCODE v24 annotations (gencode.v24.basic.annotation.gtf.gz)[37].

#### 5.1.2 Cis-regulatory elements

Candidate cis-regulatory elements (cCREs) were obtained from the ENCODE Registry of cCREs (ENCODE accession ENCSR487PRC)[18]. This dataset provides cell-agnostic cCRE annotations for GRCh38, which were used to define regulatory regions across the genome. The distribution of cCREs are shown in Figure 7 The primary cCRE file (ENCFF234XEZ.bed.gz) was downloaded from the ENCODE portal and used to filter genomic variants to those located within regulatory regions. For the ENTEx cohort, we additionally obtained donor-specific and tissue-specific cCRE annotations from ENCODE to pretrain the encoder on the chromatic activity predictions. The accession numbers for the files are provided in the Supplementary Table 2

**Figure 7.**
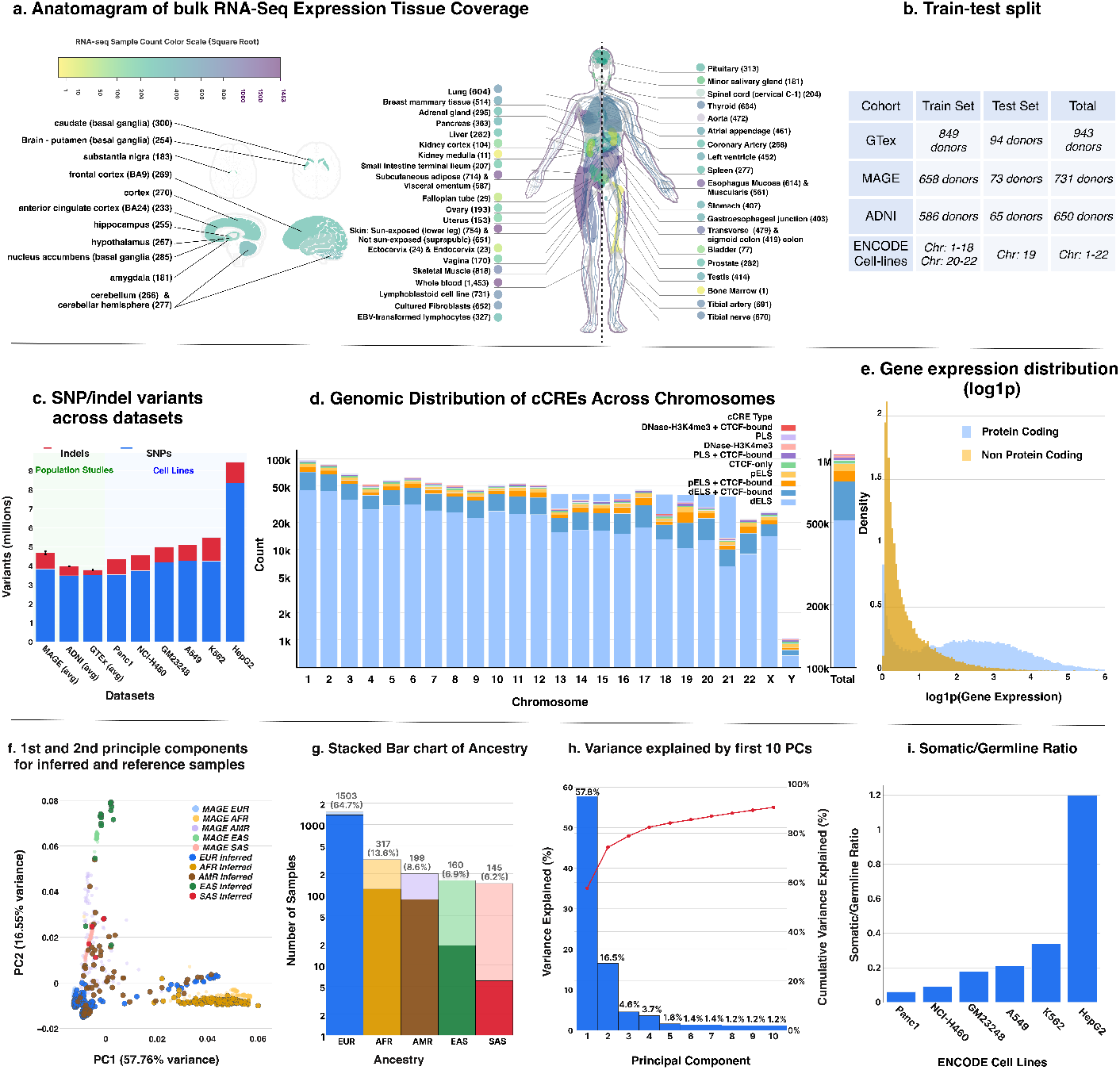
Overview of the data. **(a)** Anatomagram of RNA-seq samples over tissues. **(b)** Train-test split: the data was split by cohort donors and cell line chromosomes. **(c)** Variant distributions across population cohorts and cell lines using stacked bars (SNPs: blue; indels: red; Section S1.6). Population studies (green background) show average counts over 30 donor with error bars representing standard error of the mean (SEM): MAGE (4.7M ± 0.15M), ADNI (4.0M ± 0.08M), GTEx (3.8M ± 0.12M). Cell lines (blue background) exhibit higher total variant burden (4.3–9.4M) with HepG2 displaying maximum load. SNPs comprise ∼80–85% of variants across all datasets. **(d)** Genomic distribution of candidate *cis*-regulatory elements (cCREs) across human chromosomes (GRCh38). The right panel displays the aggregate total across all chromosomes (*n* = 1,063,878) with stacked bars representing different functional types. **(e)** Distribution of gene expression levels for protein-coding (blue) and non-protein-coding genes (yellow). Density histogram showing log-transformed gene expression values (log1p) across all gene-sample pairs. **(f)** Principal component analysis of 2,330 samples showing population structure. Light markers indicate MAGE reference populations, dark markers show inferred ancestry for GTEx and ADNI samples. **(g)** Distribution of ancestry. Light markers indicate MAGE reference populations, dark markers show inferred ancestry for GTEx and ADNI samples. **(h)** Variance explained by the first 10 principal components (bars) with cumulative variance explained (red line). **(i)** Relative somatic mutation burden in ENCODE cancer cell lines through somatic/germline variant ratios.

#### 5.1.3 Whole Genome Sequencing data

##### MAGE (1000 Genomes)

MAGE data consists of a subset of 731 donors from the 100 Genomes project. We obtained the high-coverage (30×) whole genome sequencing data from the 1000 Genomes Project [2]. Specifically, we downloaded phased SNV, INDEL, and SV variant calls for 731 samples from the 20220422 release (*1kGP_high_coverage_Illumina chrN*.*filtered*.*SNV_INDEL_SV_phased_panel*.*vcf*.*gz*, chromosomes 1-22) from the EBI FTP server (ftp.1000genomes.ebi.ac.uk).

##### ENTEx (Genotype-Tissue Expression from ENCODE)

ENTEx phased WGS data sequenced on the Illumina NextSeq 500 platform were downloaded from the ENCODE. Downloaded VCF files and their accession numbers are provided in the Supplementary Table 3.

##### GTEx v9

We obtained GTEx v9 WGS data for 953 individuals from dbGaP (study accession: phs000424.v9, genotype dataset: *phg001796*.*v1*.*GTEx_v9_WGS_953*.*genotype-calls-vcf*.*c1*.

##### ADNI (Alzheimer’s Disease Neuroimaging Initiative)

WGS data for 808 ADNI participants were obtained from the LONI Image and Data Archive. The VCF files were initially in hg19 coordinates and were lifted over to GRCh38 using CrossMap [59] with the hg19ToHg38 chain file.

##### ENCODE cancer cell-lines

Whole genome sequencing data for six ENCODE cell lines were obtained from the ENCODE Data Coordination Center: A549 (lung adenocarcinoma), GM23248 (lymphoblastoid), HepG2 (hepatocellular carcinoma), K562 (chronic myelogenous leukemia), NCI-H460 (lung large cell carcinoma), and Panc1 (pancreatic adenocarcinoma). Variant calling was performed using a dual-caller strategy combining DeepVariant v1.6.0 for germline variants and DeepSomatic v1.6.0 in tumor-only mode for somatic variant detection. DeepSomatic-identified somatic variants were prioritized at overlapping positions, with DeepVariant calls retained for non-somatic positions. All variants were filtered to retain only PASS-quality calls and annotated with variant origin (germline or somatic) using custom INFO fields. Detailed variant calling procedures and quality control metrics are provided in supplementary Section S1.4.

##### VCF filtering and processing

All WGS variant calls were filtered to retain only variants located within cCRE regions and gene transcriptional windows defined in Section 5.3.1 All the files are standardized to standard chromosome naming (adding “chr” prefix), reheadered using the GRCh38 reference, sorted, and indexed. The distributions of SNP and indels are provided in Figure 7c.

For each individual, we generated personalized DNA sequences by applying the individual’s specific variants to the reference genome sequence. We used samtools faidx to extract reference sequences and bcftools consensus to apply variants from each individual’s VCF file with haplotype consensus mode (-H I), creating allele-specific sequences for model input.

Unlike VariantFormer, Borzoi, Enformer and EVO cannot automatically handle heterozygous alternate alleles so for those models we generate the personalized sequences by considering the alternate allele using the haplotype consensus mode (-H A). For Enformer and Borzoi we consider a window of 98*kb* upstream and 98*kb* downstream from the gene start site. Whereas for EVO 2 due to computational bottleneck we considered a window of 8*kb* upstream and downstream from the gene start site.

The genotype based baseline models are fundamentally different from the sequence based model and rely on population level features to make predictions. For our genotype based models we consider a window of 150KB upstream and downstream from the gene start site and filter for variant with MAF > 0.01. The variants are further filtered to contain biallelic SNPs. The final set of variants are passed as integer genotype calls with mean imputation to the downstream random forest models.

##### Ancestry Inference

Genetic ancestry was inferred for GTEX and ADNI samples using principal component analysis and supervised classification. Linkage disequilibrium pruning was performed using PLINK v1.90b6.24 (Purcell et al. 2007) with a sliding window approach (50 SNP window, 5 SNP step, r^2^=0.2) to identify ancestry-informative markers while removing correlated variants. Principal component analysis was conducted on the LD-pruned variants, computing the first 20 principal components to capture population structure. A K-nearest neighbors classifier (k=7, distance-weighted Euclidean metric) was trained on 731 reference samples from the 1000 Genomes Project with known continental ancestry labels (African, European, East Asian, South Asian, and Admixed American). The trained classifier was applied to all GTEx and ADNI samples projected into the reference PC space to assign ancestry classifications with associated confidence scores. Detailed ancestry inference procedures, validation metrics, and concordance analyses with self-reported ethnicity are provided in Supplementary Methods S1.5.

#### 5.1.4 RNA-seq data

##### MAGE

RNA-seq data consisting of pseudo-counts for lymphoblastoid cell lines from 1000 Genomes Project participants were downloaded from Zenodo (record ID: 10535719) [49]. The data were originally annotated with GENCODE v38 gene identifiers and consisted of raw pseudo-counts. We converted the counts to TPM (Transcripts Per Million) values while considering the effective gene lengths, and normalized by per-sample read depth. Gene identifiers were subsequently mapped from GENCODE v38 to v24 using stable Ensembl gene IDs to ensure consistency across all datasets, and the data were filtered to retain only genes present in GENCODE v24.

##### GTEx v10

Gene-level TPM expression data were obtained from the GTEx Portal (*GTEx_Analysis_v10_RNASeQCv2*.*4*.*2_gene_tpm*.*gct*)[21]. To ensure high-quality data, we filtered samples with RNA Integrity Number (RIN) > 6, retaining only samples with sufficient RNA quality for reliable gene expression measurements. Expression values were standardized to GENCODE v24 annotations for consistency across all datasets.

##### ADNI

Gene expression data for ADNI participants were obtained from the Affymetrix Human Genome U219 microarray platform available through the LONI platform. We retained data only for the 650 participants who had both whole genome sequencing (WGS) and gene expression data available. Quality control filtering was performed by selecting samples with RIN > 6. For probes with multiple gene symbols, we split them into individual gene entries. Low-expression probes below the global median intensity were filtered out to reduce noise. Multiple probes mapping to the same gene were collapsed by taking the median expression value across probes. The data were then quantile-normalized to account for technical variation across arrays and log-transformed. Gene symbols were mapped to GENCODE v24 gene annotations using stable Ensembl gene IDs.

##### ENCODE cell Lines

Gene expression data for human cell lines (K562, HepG2, GM23248, A549, NCI-H460, and Panc1) were downloaded from the ENCODE Portal. We used total RNA-seq gene quantification files in TSV format with GENCODE v24 annotations aligned to GRCh38. The accession numbers are: K562 (ENCFF171FQU), HepG2 (ENCFF863QWG), GM23248 (ENCFF640FPG), A549 (ENCFF244DNJ), NCI-H460 (ENCFF876SRX), and Panc1 (ENCFF710IFD). Data were filtered to retain only genes present in the GENCODE v24 reference.

##### Data standardization and integration

RNA-seq and microarray data from all cohorts were standardized to GENCODE v24 gene annotations using stable Ensembl gene IDs to ensure consistent gene identifiers across datasets. In Figure 7 we show the rna-seq data distribution that is combined into a single AnnData framework [51]. To ensure data quality, we filtered out genes with fewer than 20 non-zero expression counts across all samples. For non-coding genes specifically, we applied stringent tissue-specific quality control by computing the 10th percentile of non-zero expression values for each tissue type (excluding cell lines) and removing expression measurements below this threshold, thereby reducing noise from low-abundance transcripts while preserving biologically relevant signals. During model training, we exclusively utilized non-zero gene expression values from the integrated dataset. The final integrated dataset contained samples profiling expression across protein-coding and non-coding genes from diverse primary tissues and cell types, enabling comprehensive modeling of gene regulatory mechanisms across the complete human genome.

#### 5.1.5 Case control data for Alzheimer’s classification

The case-control data for Alzheimer’s classification used in the preparation of this article were obtained from the Alzheimer’s Disease Neuroimaging Initiative (ADNI) database (adni.loni.usc.edu). The ADNI was launched in 2003 as a public-private partnership, led by Principal Investigator Michael W. Weiner, MD. The original goal of ADNI was to test whether serial magnetic resonance imaging (MRI), positron emission tomography (PET), other biological markers, and clinical and neuropsychological assessment can be combined to measure the progression of mild cognitive impairment (MCI) and early Alzheimer’s disease (AD). The current goals include validating biomarkers for clinical trials, improving the generalizability of ADNI data by increasing diversity in the participant cohort, and to provide data concerning the diagnosis and progression of Alzheimer’s disease to the scientific community. For up-to-date information, see adni.loni.usc.edu.

For Alzheimer’s disease (AD) classification analyses, we obtained clinical diagnosis information for ADNI participants from the ADNI Diagnosis Summary (DXSUM) data file downloaded from the LONI platform. The DXSUM file contains longitudinal diagnosis assessments for each participant across multiple study visits. For participants with multiple diagnostic assessments, we assigned the maximum diagnosis code observed across all visits to capture disease progression and ensure stable classification. Participants were classified into diagnostic categories based on standardized clinical criteria: cognitively normal controls (CN; diagnosis code = 1), mild cognitive impairment (MCI; diagnosis codes = 2), and Alzheimer’s disease (AD; diagnosis codes = 3). We ensured that the participants contain both whole genome sequencing data and gene expression measurements available. For binary AD classification, we restricted analyses to participants classified as either CN or AD, excluding MCI cases to create a clear case-control design. We created subject-level train-test splits to prevent data leakage, with 10% of subjects held out as an out-of-distribution (OOD) test set (N = 40 subjects). We additionally ensured that the OOD test data is not used during the gene expression prediction task. The remaining subjects were combined into a unified training set for model development. The final dataset comprised 330 subjects (215 AD, 155 CN) for training and evaluation across 16510 protein-coding genes in GENCODE v24.

#### 5.1.6 MAGMA preprocessing for gene enrichment analysis

To enable gene-based association analysis and quantify gene-level Alzheimer’s disease risk for model evaluation, we performed Multi-marker Analysis of GenoMic Annotation (MAGMA) v1.10 [16] preprocessing. MAGMA aggregates SNP-level associations from genome-wide association studies (GWAS) to gene-level statistics while accounting for linkage disequilibrium (LD) structure. We obtained genome-wide association summary statistics for Alzheimer’s disease from the Psychiatric Genomics Consortium (PGC) Alzheimer’s Disease Working Group [41], excluding 23andMe data due to data sharing restrictions. Summary statistics were lifted over to the GRCh38/hg38 reference genome to match GENCODE v24 gene coordinates. Duplicate SNPs at identical genomic positions were removed, retaining only the first occurrence. The 1000 Genomes Project Phase 3 European ancestry panel was used as the LD reference to account for correlation structure among variants. Using MAGMA’s annotation module, we mapped SNPs to genes based on genomic proximity. SNPs were assigned to genes if they fell within gene boundaries or within extended flanking windows of 100 kb upstream and 50 kb downstream of each gene. This window size captures regulatory variants in promoters, enhancers, and other distal regulatory elements that influence gene expression. Gene boundaries were extracted from the GENCODE v24 basic annotation, and SNP positions were obtained from the 1000 Genomes reference panel. The resulting SNP-gene annotation enabled computation of gene-level association statistics by aggregating evidence across all variants mapped to each gene while properly accounting for LD between SNPs.

#### 5.1.7 eQTL data curation

Expression quantitative trait locus (eQTL) data were obtained from the eQTL Catalogue [29] to create a comprehensive variant effect prediction benchmark. We selected six tissue-specific datasets spanning peripheral tissues from TwinsUK (adipose, blood, skin) and brain regions from Braineac2 and BrainSeq (substantia nigra, frontal cortex BA9, putamen). Summary statistics were downloaded from the eQTL Catalogue (https://www.ebi.ac.uk/eqtl/) for gene expression (ge) quantification methods. Metadata and FTP paths are provided in Supplementary Table 1.

##### Harmonization and variant selection

We perform quality control on the downloaded eQTL data such that the reference alleles are correctly mapped, and the genes has stable ENSEMBL ids in GENCODE v24. The eQTL Catalogue tissue labels were mapped to GTEx tissue nomenclature to enable direct comparison with VariantFormer predictions. For TwinsUK, tissues were harmonized as follows: “adipose” mapped to “adipose - subcutaneous”; “skin” mapped to “skin - sun exposed (lower leg)”; “blood” mapped to “whole blood”.We remove the “LCL” (lymphoblastoid cell lines) from this analysis because Borzoi does not have a lcl track. For brain studies (Braineac2 and BrainSeq), tissues were mapped as: “brain (putamen)” → “brain - putamen (basal ganglia)”; “brain (substantia nigra)” → “brain - substantia nigra”; and “brain (DLPFC)” → “brain - frontal cortex (ba9)”.

We restricted analysis to single nucleotide polymorphisms (SNPs), excluding insertions, deletions, and structural variants to maintain consistency with sequence-based predictions. For primary eQTL validation analyses, we selected the top 50,000 most statistically significant variant-gene pairs per tissue dataset, ranked by ascending p-value. This p-value-based prioritization enriches for variants with robust effect sizes while maintaining diverse allele frequencies and genomic contexts. To evaluate model performance on low-frequency variants—which are under-represented in most eQTL discovery studies but critical for precision medicine—we performed parallel analyses restricted to variants with minor allele frequency (MAF) < 0.05, selecting the top 10,000 variants per tissue within this frequency range. In addition we filtered that variant that fall withing the context windows of VariantFormer, Borzoi and Alphagenome.

##### Ancestry-specific variant stratification

We performed the population-stratified eQTL analysis using BrainSeq frontal cortex data. Variants were annotated with 1000 Genomes Project Phase super-population allele frequencies (AF_EUR, AF_AFR) obtained from pre-computed allele frequency tables. We defined two ancestry-enriched variant sets using stringent allele frequency thresholds: EUR-enriched variants required AF_EUR > 0.10 and AF_AFR < 0.05; AFR-enriched variants required AF_AFR > 0.10 and AF_EUR < 0.05. These criteria identify variants with substantial frequency differences between populations (≥ 5% frequency in one ancestry and < 5% in the other), enabling evaluation of ancestry-matched prediction accuracy. For each variant set, we retained the top 50,000 variants ranked by eQTL p-value.

The final curated datasets comprised: TwinsUK (all variants: 47,119 variant-gene-tissue combinations; MAF < 0.05: 9,604 combinations), brain tissues (all variants: 35,966 combinations; MAF < 0.05: 4,808 combinations), and ancestry-specific BrainSeq (15458 AFR/EUR stratified variant-gene pairs). All variant identifiers, genomic coordinates, eQTL effect sizes, and tissue annotations are provided in Supplementary Tables 1.

#### 5.1.8 ENCODE DNase seq data

We demonstrate that the model learns biologically meaningful gene regulatory patterns by analyzing a random subset of 50 protein-coding genes in adrenal gland tissue (UBERON:0002369). DNase-seq chromatin accessibility data for adrenal gland tissue were obtained from ENCODE (accession ENCFF632QUC) in bigWig format to serve as ground truth signals for validating model attention patterns. Gene annotations including gene start sites and strand orientation were extracted from GENCODE v24 basic annotations aligned to the GRCh38 reference genome.

#### 5.1.9 APOE genotyping data for ADNI

We extracted genotypes for APOE-defining variants from ADNI participants with AD disease status. The APOE gene contains two key single nucleotide polymorphisms (SNPs) that determine the three common alleles: rs429358 (chr19:44908684, T>C) and rs7412 (chr19:44908822, C>T). These variants define the APOE-*ε*2 (rs429358-T, rs7412-T), APOE-*ε*3 (rs429358-T, rs7412-C), and APOE-*ε*4 (rs429358-C, rs7412-C) alleles, with *ε*4 being the strongest genetic risk factor for Alzheimer’s disease. We identified AD patients from our OOD test data carrying at least one copy of risk-associated alleles (APOE-*ε*4: *N* = 12 samples with rs429358-C; APOE-*ε*2: *N* = 1 sample with rs7412-T). For each carrier, we generated two personalized genomic sequences: (1) with the individual’s actual genotype applied to the reference genome, and (2) with the APOE variant allele replaced by the reference allele (effectively converting *ε*4 or *ε*2 to *ε*3 *in silico*). This approach enabled us to quantify the risk effects of APOE variants while controlling for the individual’s genetic background across the rest of the genome.

### 5.2 Model Architecture Overview

VariantFormer employs a hierarchical two-stage architecture that integrates donor-specific genomic variants to predict personalized gene expression. The model operates on two complementary genomic contexts: cis-regulatory elements (CREs) and gene transcriptional windows, combining their representations through a multi-level transformer framework.

#### 5.2.1 Unified Mathematical Framework

Let 𝒟 = {(*d, g, t*)} represent the training data, where *d* denotes a donor with genomic variants *V*_*d*_, *g* is a target gene, and *t* is a tissue type. For each gene *g* with associated CREs ℛ_*g*_ = {*c*_1_, …, *c*_*N*_} and transcriptional window 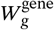, we generate donor-specific DNA sequences by embedding variants into the reference genome (detailed in Section 5.3.1).

VariantFormer models gene expression as a Poisson process, with the expression level *e*_*s*_ for sample *s* drawn from a Poisson distribution whose rate parameter is determined by the regulatory and genomic contexts. The hierarchical prediction framework can be formalized as:

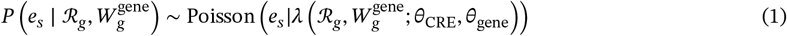

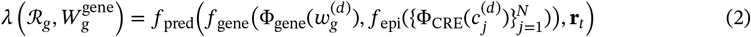

where the hierarchical architecture comprises the following components:

- Ö_CRE_: 𝒮 → ℝ^*d*^ and Φ_gene_: 𝒮 → ℝ^*d*^ – Sequence encoders (detailed in Section 5.3) that map tokenized DNA sequences to embedding space; Φ_CRE_ is frozen after pretraining while Φ_gene_ is fine-tuned
- 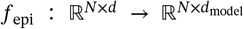 – Epigenetics Modulator capturing regulatory interactions via selfattention over CRE embeddings
- 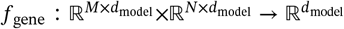 – Gene Modulator integrating gene windows with regulatory context via cross-attention
- 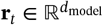 – Learnable tissue-specific registry token
- 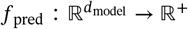 – Prediction head mapping final representation to expression level

Both *f*_epi_ and *f*_gene_ are detailed in Section 5.4. The hierarchical architecture enables multi-scale regulatory modeling through layered CRE-gene interactions:

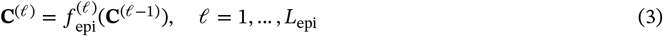

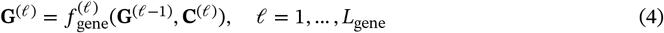

where 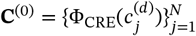 and 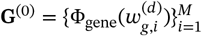 are the initial CRE and gene embeddings, respectively. This layered design mirrors biological regulatory cascades, where primary signals from local CREs are progressively integrated with higher-order combinatorial interactions.

The detailed architectural components, including BPE tokenization, encoder design, attention mechanisms, and regulatory cross-attention, are described in Sections 5.3 (pretraining) and 5.4 (gene expression prediction). Section 5.3 explains the mutation-aware pretraining strategy that grounds the model in chromatin biology, while Section 5.4 details the hierarchical modulator architecture that enables interpretable modeling of distal regulatory interactions.

### 5.3 Pretraining Sequence Encoders

#### 5.3.1 Embedding mutations into the reference genome

We constructed donor-specific, strand-aware genomic sequences by embedding small variants (SNPs and in-dels) from VCFs into the human reference (GRCh38/hg38). For each genomic region of interest (genes and their cis-regulatory elements), we first extracted the reference subsequence with samtools faidx. We then applied variants using bcftools consensus with the -H I option, which represents heterozygous genotypes using IUPAC ambiguity codes (e.g., A/G → R), and writes homozygous ALT alleles as the ALT base. Structural variants and complex rearrangements were excluded to avoid ambiguous reconstructions.

The gene coordinates are taken from GENCODE v24.

##### Per-position mutation embedding

Let *R*_chr,*p*_ denote the reference base at position *p* on chromosome chr, and let *V*_*d*_ be donor *d*’s set of small variants. The mutated base is

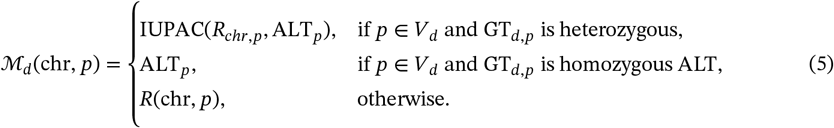

where *GT*_*d,p*_ denote the genotype of the variant. Short insertions/deletions are applied by `bcftools`; inserted bases appear at their appropriate locations in ℳ_*d*_.

##### Gene (“transcriptional”) window

For gene *i* on chromosome chr_*i*_ with start *S*_*i*_, end *E*_*i*_, and strand *o*_*i*_ ∈ {+, −}, we define the strand-agnostic window

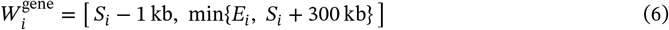

The donor-specific sequence underlying this window is

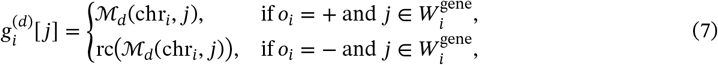

where rc(·) denotes reverse-complement. (Equivalently, build the window on the reference strand and reverse-complement the entire sequence if *σ*_*i*_ = −.)

##### Long-range cCREs and the regulatory set

To capture distal regulation, we considered a long-range window

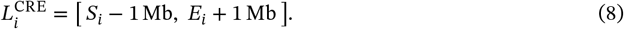

Given a dictionary of candidate cis-regulatory elements (cCREs) 𝒞 with coordinates (chr_*c*_, *a*_*c*_, *b*_*c*_), the set of elements associated with gene *i* is

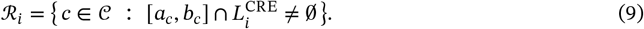

We extract a donor-specific, flanked sequence for each element *c* ∈ ℛ_*i*_ by padding δ = 50 bp on both sides:

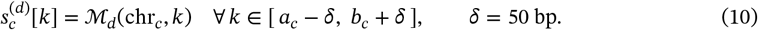

The cCREs are oriented based on the position of the gene in the forward strand vs the reverse strand.

#### 5.3.2 DNA Sequence Tokenization Framework

##### Byte-Pair Encoding Tokenizer Training on Reference cCREs

We implemented a domain-specific tokenization approach using Byte-Pair Encoding (BPE) [31] adapted for DNA sequences to effectively process genomic sequences for the downstream training. Our tokenizer is trained on candidate Cis-Regulatory Elements (cCREs) from the ENCODE project [18], specifically the comprehensive dictionary of cCRE annotations (ENCFF234XEZ) mapped to the GRCh38 human reference genome (see Section 5.1.2). For each cCRE, we extracted DNA sequences with a neighborhood extension of 50 base pairs upstream and downstream to capture potential regulatory context, resulting in variable-length sequences centered on regulatory elements.

Prior to BPE training, all DNA sequences underwent standardized preprocessing

1. **Case normalization**: All sequences were converted to uppercase for consistency.
2. **IUPAC code filtering**: We retained only valid IUPAC nucleotide codes (A, C, G, T, R, Y, S, W, K, M, B, D, H, V) and filtered out ambiguous characters.

We implemented our BPE tokenizer using the HuggingFace Tokenizers library [1], specifically employing the BPE model with a vocabulary size of 500. The tokenizer training process follows the standard BPE algorithm [31]. The trained tokenizer generates a vocabulary comprising biologically relevant DNA subsequences, ranging from individual nucleotides to longer motifs that may correspond to transcription factor binding sites or other regulatory elements. The 500-token vocabulary provides a balance between capturing sequence diversity and maintaining computational efficiency for downstream applications.

###### Algorithm 1

BPE Tokenizer Training on cCRE Sequences

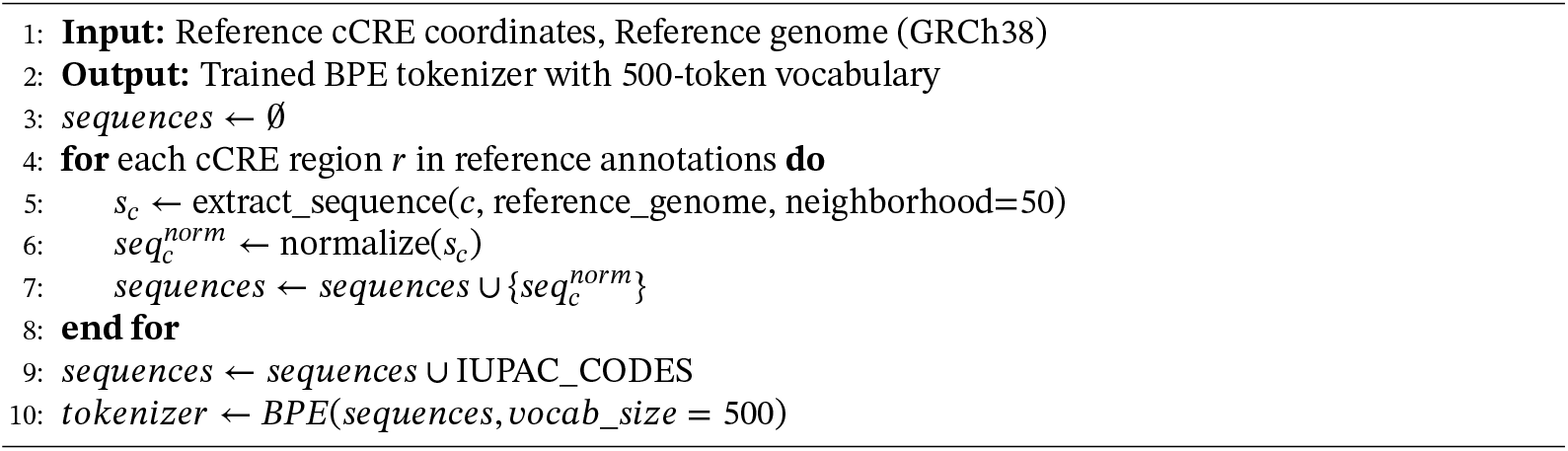

#### 5.3.3 Pretraining mutation-aware encoders on chromatin-activity data

We pretrained a mutation-aware transformer encoder (Figure 8a) to compress BPE-tokenized, donor-specific DNA sequences into regulatory representations that are sensitive to underlying variants. The encoders are trained on a auxiliary task of chromatin activity prediction task using the ENTEx data (see Section 5.1.2-5.1.3). The pre-trained encoder generates donor specific contextualized embeddings for DNA sequences that serve as the foundation for downstream gene expression prediction task.

**Figure 8.**
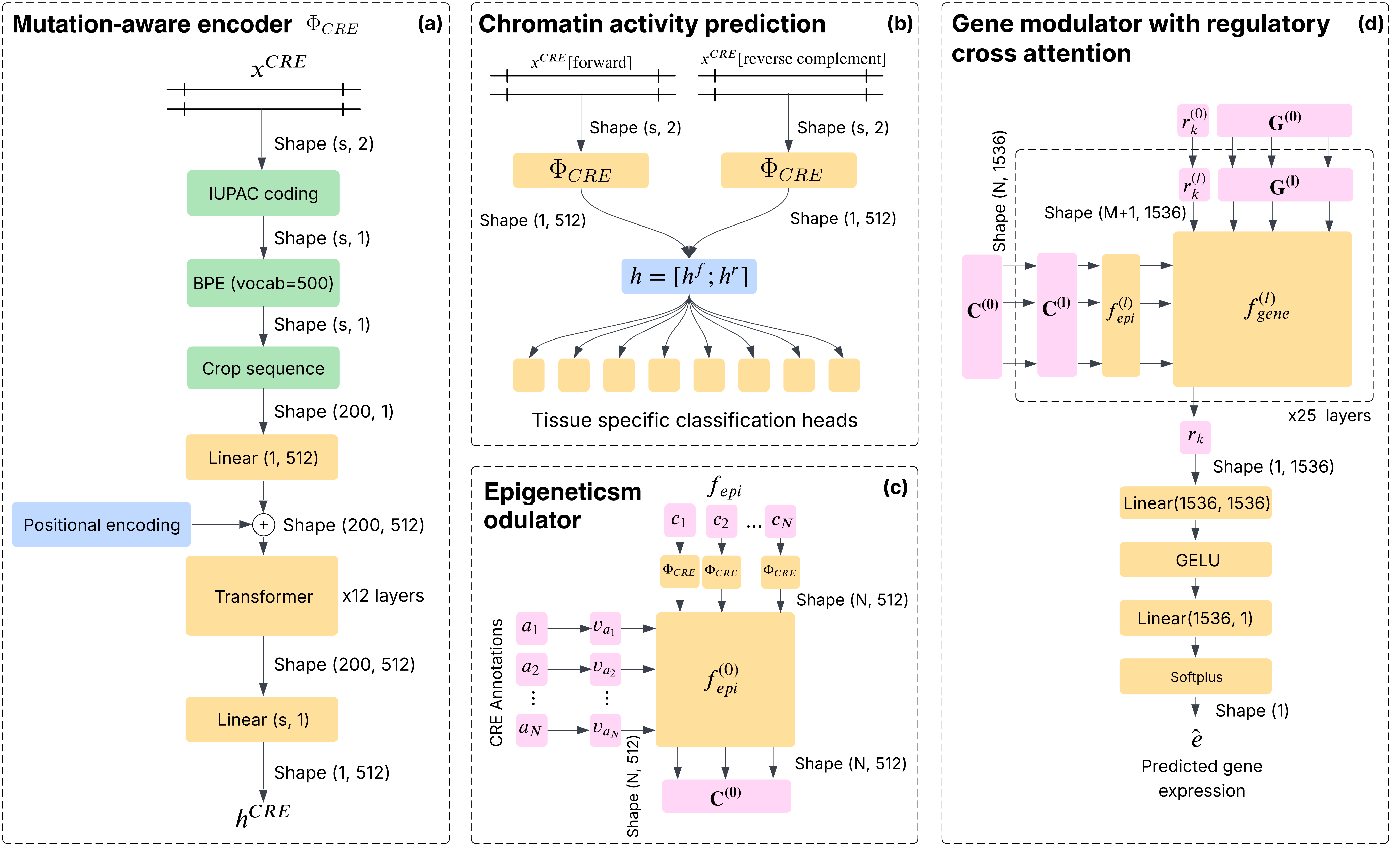
Model architecture. **(a)** Sequence encoder for pretraining on chromatin activity data. **(b)** Chromatin activity classification framework. **(c)** Epigenetics modulator *f*_*epi*_ to combine regulatory region information. CRE annotation provides contextual information about each CRE region.**(d)** Gene modulator (*f*_*gene*_ (·)) that combines sequence embedding from transcriptions window with the regulatory regions, and predict gene expression using a MLP head.

##### Architecture

The encoder maps an input token sequence *x* of length |*s*| (truncated and fixed) to a representation Φ(*x*; *θ*) ∈ ℝ^512×|*s*|^ using transformer based encoder layers consists of:

- **Token Embedding Layer**: Maps BPE tokens to a 512-dimensional embedding space
- **Positional Encoding**: ALiBi (Attention with Linear Biases) positional encoding for handling variable-length sequences
- **Multi-layer Transformer**: 12-layer transformer encoder with Flash Attention for computational efficiency
- **Tissue-specific Classification Heads**: Independent classifiers for each tissue type

The encoder processes both forward and reverse DNA strands simultaneously. We use linear pooling to combine the vector information across the sequence into a single representation vector:

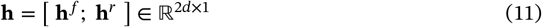

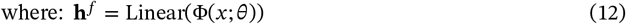

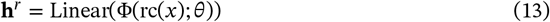

where *d* = 512. We further concatenate the representation vector coming from the forward strand and the reverse before passing through the tissue-specific classifier heads. For tissue *t* ∈*T*, a lightweight head (**W**_*t*_, **b**_*t*_) produces

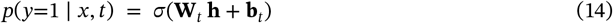

##### Classification loss

Each training sample is a tuple (*x, t, D, y*): mutated cCRE sequence *x* (with ±50 bp flank), tissue *t*, donor *D*, and binary label *y* ∈{0, 1} indicating chromatin activity. The binary labels *y* were derived by categorizing candidate regulatory elements from tissue *t* into two classes:

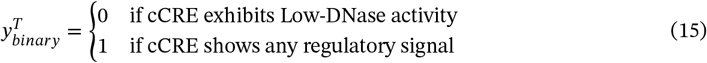

The “active” class (label 1) encompasses all cCRE categories with detectable regulatory activity, while the “inactive” class (label 0) corresponds to regions with minimal chromatin accessibility. We used per-tissue binary cross-entropy loss:

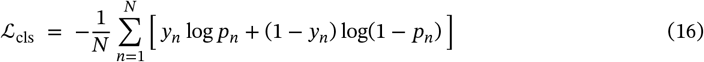

##### Mutation-aware contrastive loss

To encourage sensitivity to donor-specific sequence differences, we added a supervised contrastive loss across donors within a batch. Let 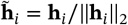 and 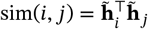. For anchor *i*, we define the negative set *A*(*i*) = {*j* ≠ *i*}. The loss is

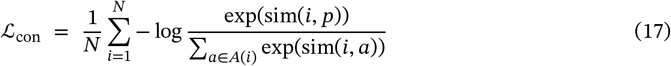

##### Total objective and batching

The total training objective combines the primary classification loss with the contrastive learning component:

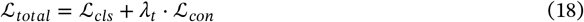

where *λ* is dynamically scheduled such that the relative magnitudes of the two loss terms remains equal. We also keep lambda bound between [.01, 100]:

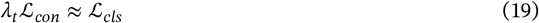

Finally, during training, we ensured that each batch contains sequences sampled uniformly from all 4 donors of the ENTEx data. This batching strategy captures effect of both tissue and mutational variability on chromatin activity.

### 5.4 Gene Expression Prediction Architecture

The gene expression prediction layer leverage the pretrained encoders to generate contextual sequence embeddings. The second level of transformer layers are trained on the the top of the embedding to prediction final gene expression level. For each gene, we extract two complementary genomic contexts: (1) cis-regulatory elements (CREs) flanking the gene within a defined neighborhood window, and (2) the transcriptional window. These regions are formally defined in Section 5.3.1.

#### 5.4.1 Transfer learning

We adapt the smutation-aware encoder for the following genomic contexts:

##### CRE Encoder (Φ_CRE_

The pretrained encoder is frozen to generate embeddings for the regulatory regions. By preserving the pretrained weights, we retain the learned representations of regulatory syntax and chromatin accessibility patterns captured during large-scale pretraining on chromatin activity data. Each CRE region 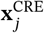 is encoded as:

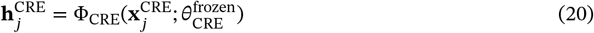

where 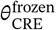 denotes the fixed pretrained parameters. This design ensures that regulatory element representations remain stable and generalizable across different prediction tasks. The sequence **x**_*j*_ is extracted from the reverse vs forward strands conditioned on the orientation of the gene.

##### Gene Encoder (Φ_gene_)

The encoder processing the transcriptional window is fine-tuned end-to-end during training. Fine-tuning enables the model to adapt the sequence representations to the specific requirements of gene expression prediction, learning features relevant to transcriptional output such as transcription factor binding motifs within gene bodies, splice site configurations, and regulatory elements in untranslated regions. The gene embedding for window *w*_*i*_ is computed as:

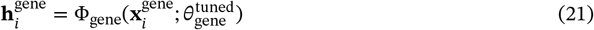

where 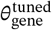 represents the fine-tuned parameters, initialized from pretraining and updated during supervised learning.

This asymmetric training strategy balances model capacity with generalization: frozen CRE representations prevent overfitting to dataset-specific regulatory patterns, while adaptive gene representations enable task-specific feature refinement.

#### 5.4.2 Sequence Partitioning and Embedding Generation

The transcriptional region is systematically partitioned into *M* = 200 non-overlapping windows of fixed length (200 tokens per window after BPE tokenization). Each window *w*_*i*_ is independently encoded through Φ_gene_ to produce a contextualized embedding 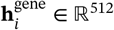, where 512 is the embedding dimension. This chunking strategy enables the model to process arbitrarily long transcriptional (spanning kilobases) while maintaining computational tractability and preserving fine-grained positional information along the gene body.

For the regulatory context, we extracted *N* CRE regions surrounding the gene, where each region is independently encoded through Φ_CRE_ to obtain embeddings 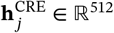.

#### 5.4.3 Projection to Unified Representation Space

To ensure dimensional compatibility across the downstream architecture, we apply learned linear projections to map both gene and CRE embeddings to a unified representation space of dimension *d*_model_ = 1536:

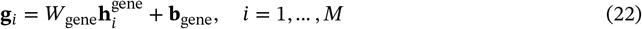

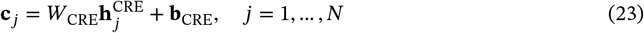

where *W*_gene_, *W*_CRE_ ∈ ℝ^1536×512^ are learnable projection matrices that serve as adaptation layers, bridging the pretrained encoder output space to the gene expression prediction architecture.

#### 5.4.4 Epigenetics Modulator for CRE Context Integration

The Epigenetics Modulator (*f*_*epi*_ (·); Figure 8c) processes the CRE embeddings 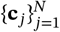 to capture long-range regulatory interactions and establish a contextualized representation of the epigenetic landscape. This module consists of *L*_E_ transformer encoder layers, each comprising:

1. **Self-attention mechanism**: Enables bidirectional communication between CRE regions to model combinatorial regulatory logic. Using Flash Attention [15] with ALiBi positional encoding [39], we compute:

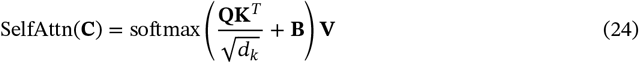

where 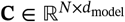 is the matrix of CRE embeddings, and **B** encodes relative positional biases. {**Q, K, V**} are obtained as linear transformation of **C**
2. **CRE annotation cross-attention**: Integrates functional annotations of CREs (e.g., PLS, dELS, pELS classifications) through a learned embedding lookup. Each CRE annotation type *a*_*j*_ is mapped to a learnable vector 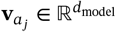, which serves as additional context:

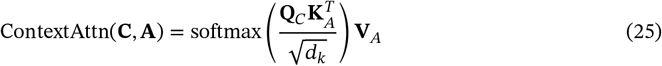

where 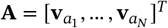 represents the annotation embeddings, {**K**_*A*_, **V**_*A*_} are linear transformation of **A**, and **Q**_*C*_ is the linear transformation of **C**.
3. **Feed-forward network**: A GeGLU activation-based MLP applies non-linear transformations:

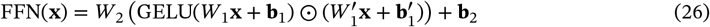
4. **Residual connection:** At each transformer layer we have a residual connection to preserve the original information and ensure gradients can flow easily during backpropagation.

The Epigenetics Modulator produces a sequence of intermediate representations 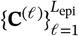, where 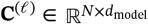 captures progressively refined regulatory context at layer *ℓ*. These multi-scale representations are subsequently utilized to modulate gene expression prediction.

#### 5.4.5 Gene Modulator with Regulatory Cross-Attention

The Gene Modulator (*f*_*gene*_(·); Figure 8d) integrates the regulatory context encoded by the Epigenetics Modulator with the intrinsic sequence information of the gene body. This module comprises *L*_G_ crossattention layers that explicitly model the regulatory influence of CREs on different regions of the gene.

For each layer *ℓ* ∈ {1, …, *L*_G_}, the gene embeddings 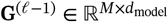 attend to the corresponding CRE representation **C**^(*ℓ*)^ from the Epigenetics Modulator:

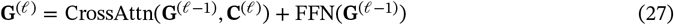

The cross-attention mechanism is defined as:

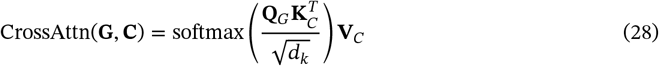

where **Q**_*G*_ = **G***W*_*Q*_, **K**_*C*_ = **C***W*_*K*_, and **V**_*C*_ = **C***W*_*V*_ with learnable projection matrices. This formulation allows each gene window to selectively integrate information from relevant regulatory regions, capturing mechanisms such as enhancer-promoter looping and distal regulatory effects.

#### 5.4.6 Conditional Tokens for Tissue-Specific Contextualization

To enable tissue-specific gene expression prediction, we introduce learnable conditional tokens that capture cell-line and tissue-specific regulatory programs.

##### Multi-registry tokens

We employ a tissue and cell-line specific registry tokens 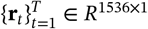, where *T* is the number of tissues and *d*_*model*_ = 1536. For a sample from tissue *t*, the corresponding registry token **r**_*t*_ is prepended:

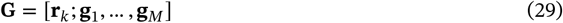

where [**g**_*i*_, …, **g**_*M*_] correspond to the transcription window for that specific sample. This enables the model to learn tissue-specific aggregation strategies and offer greater flexibility for multi-tissue learning.

#### 5.4.7 Hierarchical CRE-Gene Interactions

A key architectural innovation is the hierarchical interaction between CRE and gene regions across multiple transformer layers. Unlike conventional approaches that aggregate regulatory information in a single step, our model establishes a layered communication pathway:

1. **Layer 1**: The Gene Modulator attends to the initial CRE representation **C**^(1)^, capturing primary regulatory signals (e.g., promoter accessibility, nearby enhancer activity).
2. **Layer 2 to** *L*_**gene**_: Each subsequent layer attends to progressively refined CRE representations **C**^(*ℓ*)^. This hierarchy allows the model to first capture local regulatory effects, then integrate longer-range interactions and combinatorial logic as encoded by deeper Epigenetics Modulator layers.

This design mirrors biological regulatory cascades, where primary regulatory signals are established by local CREs, and higher-order interactions emerge through cooperative binding and chromatin remodeling.

The attention patterns in the cross-attention layers provide mechanistic insights into regulatory logic. High attention weights between a gene window and specific CREs indicate predicted regulatory influence, enabling interpretation of model predictions in terms of known or putative regulatory interactions.

#### 5.4.8 Expression Prediction Head

The final gene representation, obtained by pooling the output of the Gene Modulator, is passed to a shared prediction head that operates across all tissues. The pooled representation 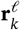 (the tissue specific registry token embedding) is transformed through a two-layer MLP:

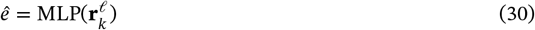

where *h* is a shared MLP defined as:

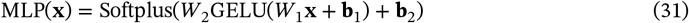

The use of a single shared prediction head promotes parameter efficiency, and the tissue-specific variation is captured through the conditional tokens. The softplus activation ensures non-negative predictions compatible with Poisson negative log-likelihood loss, which appropriately models the count-based nature of RNA-seq data:

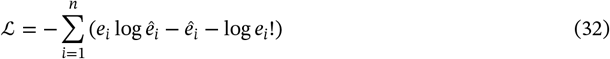

where *e*_*i*_ and 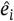 are the log 1*p* normalized observed and predicted expression levels (TPM) for sample *i*. The Poisson distribution naturally accounts for the non-negative nature of transcript counts inherent in sequencing data. We approximate *e*! with Stirling’s approximation to account for the non-integer nature of the log normalized count data.

### 5.5 Training procedure

#### Pretraining of the encoders (Φ_*CRE*_, Φ_*gene*_)

The encoder pretraining was performed using distributed data parallel training across 8 GPUs with a per-device batch size of 32 and gradient accumulation over 12 steps, yielding an effective batch size of 3,072 sequences. Training proceeded for 29 epochs using the Adam optimizer with an initial learning rate of 10^−4^ and weight decay of 0.01. Learning rate scheduling employed ReduceLROnPlateau with a patience of 2 epochs and reduction factor of 0.1, monitoring validation loss to adaptively scale the learning rate when performance plateaued. The model converged after 29 epochs, at which point validation loss stabilized. All computations utilized mixed-precision training (bfloat16) to accelerate training and reduce memory consumption while maintaining numerical stability.

#### Pretraining of gene expression prediction modules (Φ_*gene*_, *f*_*epi*_, *f*_*gene*_)

Gene expression prediction training was conducted in two sequential stages to progressively expand model capacity and gene coverage.

#### Stage 1 (Protein-coding genes)

Initial training focused on 18,439 protein-coding genes over 12 epochs. The CRE encoder remained frozen (Φ_CRE_ in evaluation mode) to preserve pretrained regulatory representations, while the gene encoder (Φ_*gene*_) was finetuned, and the modulators (*f*_*epi*_, *f*_*gene*_) were trained end-to-end. Training employed distributed data parallel across 376 GPUs with per-device batch size of 2 gene-donor pairs and gradient accumulation over 11 steps. Since each gene-donor pair is expanded to 2 tissue-specific samples through weighted tissue sampling, the effective batch size was 16,544 samples (376 GPUs × 2 pairs × 2 tissues × 11 accumulation steps). Optimization used AdamW with initial learning rate 10^−4^, weight decay 0.01, and gradient clipping at 1.0. Learning rate scheduling followed a warmup-cosine strategy with 1% warmup steps followed by cosine annealing to minimum learning rate 10^−5^. Mixed-precision training (bfloat16) accelerated computation while maintaining numerical stability.

#### Stage 2 (All annotated genes)

Training continued for 10 additional epochs incorporating 32,517 non-coding genes (50,956 total genes), initializing from the parameters of Stage 1. The learning rate was reduced to 4 × 10^−5^ with minimum learning rate 10^−6^ to enable fine-grained adaptation without disrupting learned protein-coding representations.

### 5.6 Train-test split for model pretraining

In phase 1 we pretrained the mutation-aware encoders on ENTEx paired whole genome sequencing (WGS) and cis-regulatory element (cCRE) activity data. The cCRE activity profiles were obtained from 16 tissues across four ENTEx donors (ENCDO271OUW, ENCDO451RUA, ENCDO793LXB, and ENCDO845WKR). The tissue types and corresponding ENCODE accession identifiers are provided in Supplementary Table 2. For encoder pretraining, we used chromosomes 1-22 excluding chromosome 21 for training. Chromosome 21 was designated as the held-out test chromosome to evaluate the model’s ability to predict chromatin accessibility patterns in unseen genomic regions.

During phase 2 we created held-out test sets ensuring no subject-level data leakage between training and test partitions. For the MAGE dataset, we stratified subjects by superpopulation ancestry (African, American, East Asian, European, and South Asian from the 1000 Genomes Project) and randomly sampled 10% of subjects from each population stratum for the test set. This stratified sampling ensured balanced representation of genetic diversity in the evaluation. For GTEx and ADNI datasets, we randomly sampled 10% of unique subjects for the test sets, with all tissue samples from a given subject assigned exclusively to either the training or test partition to prevent data leakage. For the ENCODE cell line data, we employed a chromosome-based holdout strategy, designating all gene expression measurements on chromosome 19 as the test set while all other chromosomes constituted the training set. Figure 7 illustrates the distribution of samples across training and test partitions for all cohorts.

## 6 Post training Tasks

### 6.1 Gene expression prediction task

#### 6.1.1 Baselines

We compared the performance of VariantFormer, against several published SOTA models for linking genetic variation to gene expression. Benchmark models were selected to span the major methodological paradigms: statistical genetics approaches (genotype-RF; Section S1.1), sequence-based deep learning frameworks (Borzoi, Enformer; Section S1.2), and evolutionary-contextual models (EVO; Section S1.2). This strategy ensures that comparisons highlight both incremental and orthogonal advances introduced by VariantFormer. We have not benchmarked AlphaGenome [5] as the model was not released for downstream fine tuning.

#### 6.1.2 Correlation metrics for performance quantification

We evaluate gene expression prediction accuracy using gene correlation and subject correlation. These metrics provide distinct perspectives on prediction quality by measuring variability across different axes of the data.

##### Gene correlation

Gene correlation quantifies how accurately the model predicts expression variability across samples for individual genes. For each gene *g*, let 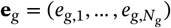 denote the vector of observed expression values and 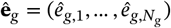 the corresponding model predictions across all *N*_*g*_ samples. The gene correlation is computed as:

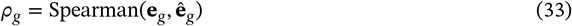

where Spearman denotes the Spearman rank correlation coefficient. This metric captures the model’s ability to predict individual-level and tissue-specific variation for each gene.

##### Subject correlation

Subject correlation quantifies how accurately the model predicts expression variability across genes for individual samples. For each sample *s* (defined by a unique donor-tissue combination), let 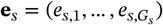 denote the vector of observed expression values and 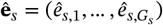 the corresponding predictions across all *G*_*s*_ genes profiled in that sample. The subject-level correlation is:

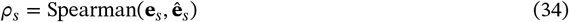

This metric primarily reflects the model’s ability to capture sample-level performance across genes.

#### 6.1.3 Ancestry-specific gene identification

We applied one-way ANOVA to identify the ancestry-specific genes that vary significantly across genetic ancestries. of the test data coming from MAGE.

For each gene *g* in the held-out test set, we aggregated expression measurements 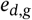 across all donors *d* within each ancestry group, yielding five population-specific expression distributions. We then performed one-way ANOVA to test the null hypothesis that mean expression levels are equal across all five ancestries:

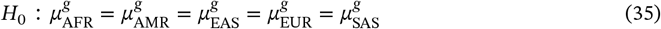

where 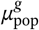 denotes the population mean expression for gene *g* in ancestry group pop. To control for multiple testing across all genes, we applied a stringent Bonferroni-corrected significance threshold of *p* < 10^−5^. Genes satisfying this threshold were designated as ancestry-specific and retained for downstream performance evaluation.

### 6.2 Alzheimer’s disease risk analysis

#### 6.2.1 Zero-shot scoring strategy in Alzheimer’s disease risk analysis

We assess VariantFormer’s embedding geometry to capture genetically grounded Alzheimer’s disease (AD) signal using MAGMA’s gene enrichment analysis [16]. For each gene *g*, tissue *t*, and donor *d*, the model outputs an embedding vector **r**_*t*_ (*g, d*) summarizing donor-specific information in tissue *t*.

##### Embedding based zeroshot scores

From cognitively normal (CN) donors 𝒟_0_ we compute a CN centroid

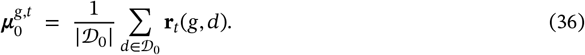

We then define a *gene-level* separability scores by averaging distances from this centroid across donors in the analysis cohort 𝒟:

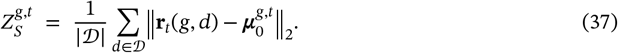

These 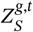 values are continuous gene properties supplied to MAGMA as covariates.

##### MAGMA gene-property regression

MAGMA aggregates SNP-level GWAS results into per-gene association *p*-values *p*_*g*_ and corresponding *Z*-scores 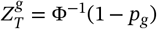. For each tissue *k*, MAGMA fits a gene-level generalized least square model:

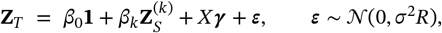

where 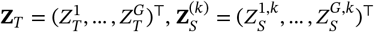, *X* includes gene-level controls (e.g., size, density, logs), and *R* is the LD-informed gene–gene correlation matrix. We test *H*_0_: *β*_*k*_ = 0 versus *H*_*A*_: *β*_*k*_ > 0 and report the corresponding one-sided *p*-value. This LD-aware regression calibrates embedding separability against inherited risk while account for the LD structure and gene properties.

This zero-shot analysis evaluates whether VariantFormer’s learned representations encode heritable AD risk without any training on disease phenotypes. Critically, the embedding-derived separability scores are computed from sequence-based gene representations alone, with no knowledge of AD status during model training or embedding generation. A statistically significant positive *β*_*k*_ indicates that genes whose embeddings discriminate AD from CN donors in our small ADNI cohort (*N* = 370) are enriched for the same genes that show association with AD in large-scale GWAS meta-analyses (*N* > 100,000 individuals). This enrichment validates that VariantFormer has learned biologically meaningful regulatory representations that capture population-level genetic architecture of disease risk, rather than merely fitting to the training data.

#### 6.2.2 Supervised classification framework in Alzheimer’s disease risk analysis

We perform binary classification of AD status from sequence-model embeddings. Let *G, D*, and *T* denote the sets of genes, donors, and tissues, respectively. For each *g* ∈ *G* and *d* ∈ *D* we observe a label *y*_*g,d*_ ∈ {0, 1} (control vs. AD). For every gene *g*, donors are partitioned once into a *training* set 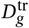 and a (OOD) test set 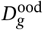 (Section 5.1.5; this split is fixed and reused for all models. We ensure that there is no overlap between the donors in the train and test set.

##### Embedding maps

Each sequence model *m* defines a feature map:

- **Tissue-agnostic models** (*m* ∈ {Enformer, Borzoi, EVO 2, genotype-RF}): 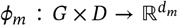 yields 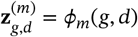.
- **VariantFormer** (gene×tissue): 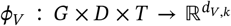 yields 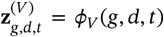 by selecting exactly the embedding coordinates annotated with tissue *t*.

We leverage the same gene embeddings for the baseline models as used in the gene expression prediction (Section S1.2). For this particular dataset, we used a total of 16,510 protein coding genes.

##### Classifier

We train random forest classifiers to ppredict the disease label. For tissue-agnostic models (per gene) we build a random forest model for each gene, in comparison for VariantFormer (per gene–tissue pair) we trained a random forest model for each gene-tissue pair. We performed a grid search to find the optimal number of trees ∈ {100, 500, 1000} and maximum depth ∈ {10, 20, 30}).

##### Cross-validation and model selection

For each modeling unit (a gene *g* for tissue-agnostic models; a gene–tissue pair (*g, k*) for VariantFormer), we run *stratified K*-fold (*K*=10) cross-validation on the training donors, preserving case/control balance per fold. We pick the best hyperparameter based on cross-validation results and then refit once on all training donors with the selected setting and compute OOD probabilities on 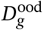.

##### Training-only Top-*K* selection

To avoid test leakage when aggregating performance, we use only the cross validation predictions to rank units by AUROC and retain the Top-*K* for reporting. For tissue-agnostic models we rank genes and keep the Top-*K* genes; for VariantFormer we rank gene–tissue pairs (*g, k*) and keep the Top-*K* pairs. All final summaries are computed solely from the OOD predictions of these selected units. For VariantFormer, we treat each (*d, k*) as a distinct observational unit for aggregation and uncertainty.

##### Metrics and aggregation

For each selected unit we compute AUPRC on its OOD scores, and we report the average across top-k selected units. In a similar fashion we report the cross-validation scores.

##### Uncertainty estimation

We quantify uncertainty with a *nested bootstrap*. In each of *B* = 2000 replicates, we resample donors with replacement within each selected unit, recompute that unit’s AUROC/AUPRC, then resample units with replacement across the selected set and average to obtain a replicate-level estimate. We report 95% percentile intervals over the *B* replicates.

##### Comparative evaluation

We apply the same protocol to VariantFormer, Enformer, Borzoi, EVO 2, and Genotype. The only structural difference is the definition of a modeling unit (gene–tissue for VariantFormer; gene for the others). We report mean AUPRC on gene-specific OOD donors with nested-bootstrap 95% confidence intervals.

### 6.3 Variant based analysis

#### 6.3.1 eQTL scoring strategy

We developed a population-aware scoring framework that leverages VariantFormer’s ability to generate expression predictions from personalized genome sequences. For each variant-gene-tissue combination, we generated two sets of predictions: (i) a reference baseline using the GRCh38 reference genome sequence, and (ii) ancestry-matched predictions where the focal variant was introduced into population-specific genomic contexts of the five 1000 Genomes super-populations (AFR, EUR, EAS, SAS, AMR).

Let 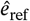 denote the predicted expression from the reference genome and 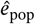 the prediction from the population-specific genome with the variant allele. The variant effect score for population *p* is computed as:

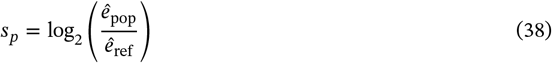

This log_2_ fold-change formulation provides an interpretable effect size metric analogous to eQTL beta coefficients, with positive values indicating expression increases and negative values indicating decreases. Additionally, to compute a single variant effect score that accounts for global allele frequency distributions we generated an allele-frequency–weighted ensemble across the five populations:

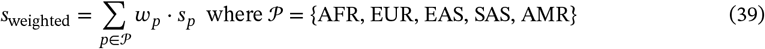

where weights 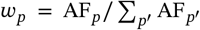 are derived from 1000 Genomes Phase 3 super-population allele frequencies. This weighted average prioritizes predictions from populations where the variant is more common, providing a frequency-calibrated estimate of average regulatory impact.

#### 6.3.2 In silico mutagenesis of APOE alleles

We performed in silico mutagenesis on APOE variants—the strongest genetic risk factors for Alzheimer’s disease to evaluate VariantFormer’s capacity to predict variant-attributable disease risk through counterfactual genome editing. For each carrier, we generated VariantFormer predictions under two genomic scenarios: (i) the patient’s observed genotype with the APOE variant present, and (ii) an in silico edited genome where the variant allele was computationally reverted to the reference allele (effectively converting *ε*4 or *ε*2 to the neutral *ε*3 allele). For each genomic scenario (observed and edited), VariantFormer generated APOE gene embeddings **r**_*k*_ ∈ ℝ^*d*^ across *T* = 13 brain tissues. These embeddings were converted to tissue-specific AD risk probabilities *p*_*k*_ ∈ [0, 1] using random forest classifiers trained on the Alzheimer’s disease classification task (Section 6.2.2). Each classifier was trained to discriminate AD versus cognitively normal (CN) donors based on gene embeddings, enabling conversion of sequence-derived representations into disease risk predictions.

##### Log odds ratio calculation

We computed log odds ratios comparing AD risk under the observed variant genotype versus the in silico edited background. For each tissue *k* and donor *d*, odds were calculated as odds_*k,d*_ = *p*_*k,d*_/(1 − *p*_*k,d*_). The log odds ratio for donor *d* was then:

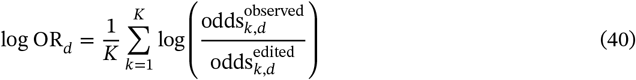

aggregating evidence across all brain tissues. Positive log odds ratios indicate that the variant increases AD risk relative to the neutral allele; negative values indicate protective effects. We report mean log odds ratios across all carriers of each variant type, with 95% confidence intervals computed using the standard error of the mean.

This counterfactual framework provides a computational approach to dissect variant contributions to complex disease phenotypes without requiring matched experimental perturbations, enabling evaluation of whether VariantFormer’s predictions align with established genetic architecture (APOE-*ε*4 risk-increasing, APOE-*ε*2 protective).

### 6.4 Cross domain analysis

#### 6.4.1 Correlation between attention and DNAse-seq

To assess whether VariantFormer’s learned attention patterns capture biologically meaningful regulatory information, we evaluated concordance between model attention weights and experimentally measured chromatin accessibility. We analyzed 50 randomly selected protein-coding genes from GENCODE v24 using reference genome sequences conditioned on adrenal gland tissue context.

VariantFormer employs 25 gene modulator layers, each with 32 attention heads (800 layer-head combinations total). For each gene, we extracted attention weights between the tissue-specific registry token and all CRE embeddings: 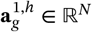, where *N* is the number of cCREs associated with the gene. These attention weights represent the importance assigned by the model to each regulatory element when predicting gene expression. For each layer-head pair, we computed Spearman correlation between attention scores across cCREs and corresponding DNase-seq signal intensities (ENCODE accession ENCFF632QUC) extracted from bigWig files at CRE coordinates using `pyBigWig`. The layer-head pair achieving the highest Spearman correlation was designated as optimal for that gene. Finally, for visualization, the cre-based attention scores were converted to bigWig format as shown in Figure 6.

#### 6.4.2 Gene representation using VariantFormer embeddings

We generated gene-level embeddings using VariantFormer-AG conditioned on reference genome sequences for two tissues: brain-cortex and whole blood. For each gene *g* among all genes, we extracted tissue-specific embeddings denoted as 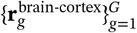 and 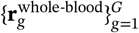.

To visualize the tissue-specific organization of the learned latent space, we performed two complementary analyses. First, embeddings from both tissues were concatenated and jointly projected into two-dimensional space using UMAP to assess tissue separability. Second, for more granular exploration of within-tissue structure, we separately projected embeddings from each tissue using tissue-specific UMAP. Prior to running UMAP for all visualizations, embedding dimensionality was reduced from 1536 to 50 using PCA to improve computational efficiency and reduce noise.

## 7 Data availability

The datasets analyzed in this study are subject to controlled access agreements. GTEx and ADNI data, including all processed derivatives, are available through managed access applications only and cannot be shared directly. Researchers interested in accessing these datasets must apply through the appropriate data access committees. Original data sources:

1. GTEx controlled access data: Apply through dbGaP (phs000424)
2. ADNI controlled access data: Apply at http://adni.loni.usc.edu/data-samples/access-data/
3. ENCODE open access data: https://www.encodeproject.org/ (publicly available)
4. 1000 Genomes Project open access data: https://www.internationalgenome.org/ (publicly available)

Due to data use agreement restrictions, processed GTEx and ADNI data files cannot be shared. Researchers must obtain approval from the respective data access committees and process the data independently using our provided analysis pipeline.

## 8 Code availability

The pretrained models are publicly available on GitHub https://github.com/czi-ai/variantformer and on CZI’s virtual cells platform https://virtualcellmodels.cziscience.com/model/variantformer.

## 9 Acknowledgments

We thank Maximilian Lombardo for introducing the browser-based user experience for model interaction on the Virtual Cell Platform and creating narrative videos that demonstrate the model’s impact; Ambrose Carr and Eric Wang for assisting with data access; Wyatt Robarts, Jae Lee, and Miguel Molina for legal and security counsel; Krunal Patel, Sanchit Gupta and CZI AI engineering team for providing research engineering support, Manu Mishra, McCullen Sandora and CZI AI Infra team for supporting the computing needs; Yun Liu and the CZI Creative team for assistance with figure preparation and graphic design; Timmy Huang, Steve Herrin, Manasa Venkatakrishnan and Omar Valenzuela for assisting with Notebook tutorials and open sourcing the code repository; Qing Mao for her valuable contribution to benchmarking; and Amanda Mahoney for helpful guidance on communications and outreach related to this preprint.

Funding for ADNI: Data collection and sharing for the Alzheimer’s Disease Neuroimaging Initiative (ADNI) is funded by the National Institute on Aging (National Institutes of Health Grant U19AG024904). The grantee organization is the Northern California Institute for Research and Education. In the past, ADNI has also received funding from the National Institute of Biomedical Imaging and Bioengineering, the Canadian Institutes of Health Research, and private sector contributions through the Foundation for the National Institutes of Health (FNIH) including generous contributions from the following: AbbVie, Alzheimer’s Association; Alzheimer’s Drug Discovery Foundation; Araclon Biotech; BioClinica, Inc.; Biogen; Bristol-Myers Squibb Company; CereSpir, Inc.; Cogstate; Eisai Inc.; Elan Pharmaceuticals, Inc.; Eli Lilly and Company; EuroImmun; F. Hoffmann-La Roche Ltd and its affiliated company Genentech, Inc.; Fujirebio; GE Healthcare; IXICO Ltd.; Janssen Alzheimer Immunotherapy Research & Development, LLC.; Johnson & Johnson Pharmaceutical Research & Development LLC.; Lumosity; Lundbeck; Merck & Co., Inc.; Meso Scale Diagnostics, LLC.; NeuroRx Research; Neurotrack Technologies; Novartis Pharmaceuticals Corporation; Pfizer Inc.; Piramal Imaging; Servier; Takeda Pharmaceutical Company; and Transition Therapeutics.

## S1 Supplementary Information

### S1.1 Gene expression prediction with genotype based random forest model

#### Problem setup

We model tissue-specific gene expression as a function of local genotype. For each gene *g*, tissue *t*, and donor *d*, we denote the observed expression by *e*_*d,t,g*_ ∈ ℝ. For every gene *g* we train an independent random-forest regressor to predict *e*_*d,t,g*_ from variant dosages in a ±150 kb cis-regulatory window around *g*, allowing nonlinear relationships between genotype and expression across donors and tissues.

Let *n*_*d*_ be the number of donors for gene *g, p*_*g*_ the number of variants in the cis-window, and *T* the number of tissues. We first form a genotype matrix 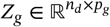 (rows: donors; columns: variants). After appending a one-hot vector of tissue identity, the design matrix for a gene–tissue pair (*g, t*) is 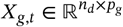, and the target vector is 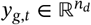.

##### Gene expression labels

We extract expression from the processed RNA-seq matrix and remove zero-valued samples. We then log-transform to stabilize variance:

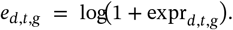

##### Genomic window

Each gene *g* is mapped to chromosome chr(*g*), strand, and GENCODE coordinates start(*g*), end(*g*). We define

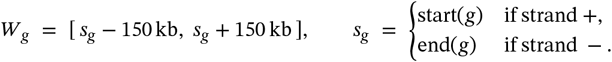

##### Genotype matrix construction

We extract all SNVs within *W*_*g*_ from the cohort VCF using bcftools [14]. For donor *d* and variant *v*, the allele counts *a*_1_, *a*_2_ ∈ {0, 1} (0 = reference, 1 = alternate) define the dosage

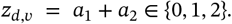

Variants with missing rate *m*_*v*_ > 0.99 are discarded. Remaining missing values are mean-imputed:

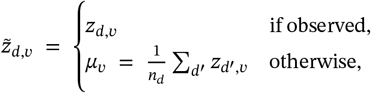

with *μ*_*v*_ = 0 if all genotypes at *v* are missing. We set 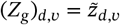.

##### Model training

For each (*g, t*) we fit a GPU-accelerated cuML random-forest regressor. We select hyperparameters (number of trees, maximum depth, minimum samples split) by 3-fold cross-validation.

To summarize for every gene–tissue pair (*g, t*) we use genotypes in a ±150 kb cis-window to build *Z*_*g*_ and train a random-forest regressor to predict log-transformed expression *e*_*g,k*_ across donors *d* and tissues *k*. This captures genotype and tissue-specific regulatory effects within a unified, nonlinear model.

### S1.2 Finetuning DNA based models (Borzoi, Enformer, and EVO 2) for gene expression prediction

For DNA based baselines we selected: Enformer [4], Borzoi [6] and EVO 2 [9].

#### Enformer

A transformer based architecture with a 196 kb receptive field trained to predict multiple genomic tracks—including chromatin accessibility, transcription factor binding, histone modifications, and CAGE-based expression—from sequence. Enformer is trained on the reference genome human and mouse without taking into account personal mutations.

#### Borzoi

Similar to Enformer, Borzoi is a convolutional–transformer architecture trained to predict tissuespecific RNA-seq coverage directly from DNA sequence. Borzoi processes genomic sequences up to 524 kb, producing high-resolution predictions of RNA abundance across diverse cell and tissue types. For comparability with Enformer, we limited Borzoi’s receptive field to the 196 kb context window used in Enformer. Borzoi was trained on bulk human and mouse data, including RNA-seq from GTEx and ENCODE, as well as complementary functional assays such as DNase, ChIP-seq, ATAC-seq, and CAGE.

#### EVO 2

An evolutionary-contextual foundation model trained in a zero-shot, self-supervised manner on 9.3 trillion DNA base pairs spanning all domains of life, with parameter scales up to 40B and a receptive field of 1 million base pairs. We however restricted the context window to 8192 kb due to GPU memory limitations. Unlike Enformer and Borzoi, which are trained on functional genomics assays, EVO 2 learns regulatory grammar directly from raw sequence and comparative evolutionary constraint via an auto-regressive self-supervision, enabling it to identify biologically meaningful features such as exon–intron boundaries and transcription factor binding motifs. Because EVO 2 is grounded in aggregate evolutionary conservation and does not incorporate personalized genomes or tissue-specific expression context, benchmarking against it highlights whether Variantformer’s genotype-aware and tissue-context embeddings confer predictive advantages beyond conservation-based inference.

For Enformer and Borzoi, we use a center pooling to accumulate the sequence level embedding which is later used for the downstream finetuning task. For EVO 2, we used the embedding from the last token. Table S1 specifies model parameters that were used to create the embeddings for each gene and donor pair.

**Table S1.** Summary of models benchmarked against VariantFormer.

| Model | Parameters | Context | Emb. Dim. | Embedding Strategy |
| --- | --- | --- | --- | --- |
| Enformer | ~500M | ~196 kb | 3072 | Center pooling (10 tokens) |
| Borzoi | ~600M | ~196 kb | 1536 | Center pooling (10 tokens) |
| EVO 2 | ~7B | ~8 kb | 4096 | Last-token embedding |

#### Finetuning strategy

To ensure comparability across benchmarks, we trained tissue-specific multilayer perceptron (MLP) heads on top of the pretrained DNA embeddings produced by Enformer, Borzoi, and EVO 2. This fine-tuning procedure enabled each model to generate gene-level expression predictions within a given tissue context, thereby aligning them with the donor-level prediction setting of VariantFormer. In addition to expression prediction, this framework allowed systematic evaluation of each model’s ability to capture the effects of variants on personalized genomes, highlighting the relative advantages of VariantFormer’s genotype-aware embeddings over models trained exclusively on reference or evolutionary sequence data.

#### Model Architecture

The tissue-specific prediction architecture consists of a shared linear projection layer that maps each baseline model’s embeddings to a common dimensional space, followed by dedicated multi-layer perceptron (MLP) networks for each of the 63 tissues. Each tissue-specific MLP contains three linear layers with layer normalization, GELU activations, and dropout regularization, culminating in a single output neuron that predicts gene expression levels. Training employs a subsampling strategy where tissues are selected per gene-donor pair in each batch, with complete tissue coverage achieved over the full training cycle.

#### Training configuration

Baseline Models were finetuned on 64 H100 GPUs, with a per-GPU batch size of 8 gene-donor pairs. In each batch 2 tissues are subsampled and gradients are accumulated over 24 batches, resulting in an effective batch size of 24576, for all three foundation models. The learning rate warms up to 1e-4 over the first 1% of training steps, then follows cosine annealing decay to 1e-5, with 0.01 weight decay applied. We trained two models for each baseline: one with ∼ 18*K* genes which are protein coding and one with ∼ 50*k* genes that encompass other non protein coding genes. The model for protein coding genes was trained to 15 epochs. This model was trained for an additional 10 epochs on a larger gene set similar to the training paradigm of transcriptformer.

### S1.3 Baseline Variant Scoring Strategies

#### S1.3.1 AlphaGenome Variant Scoring

AlphaGenome [5] is a large-scale genomic foundation model that predicts the functional effects of genetic variants on molecular phenotypes. For our variant effect prediction benchmark, we utilized AlphaGenome’s RNA-seq scoring capability through their API.

##### Prediction Pipeline

The AlphaGenome scoring pipeline processes variants through the following steps:

1. **Variant Filtering:** Input variants are filtered to remove entries containing ambiguous bases (N) in either the reference or alternate allele. Variants must include standard genomic coordinates (chromosome, position) and alleles (REF, ALT).
2. **Sequence Context:** For each variant, a genomic interval is constructed around the variant position. We used a sequence length of 1 MB (configurable from 2 KB to 1 MB) to provide sufficient context for the model to capture long-range regulatory interactions.
3. **Tissue-Specific Scoring:** AlphaGenome predicts tissue-specific effects by leveraging GTEx tissue annotations. Our pipeline maps tissue identifiers from our dataset vocabulary to AlphaGenome’s tissue keys using a predefined mapping file.
4. **Gene-Specific Predictions:** For each variant-tissue pair, AlphaGenome generates predictions for all genes within the genomic interval. We extract scores specific to the target gene by filtering on gene ID.
5. **Score Aggregation:** The raw scores from AlphaGenome represent predicted changes in RNA-seq expression levels. For genes with multiple isoforms or prediction windows, we compute the mean raw score across all predictions.

##### Model Configuration

We used the following AlphaGenome configuration:

- Organism: Homo sapiens (human)
- Output type: RNA-seq predictions
- Sequence length: 1 MB context window
- Scorer: RECOMMENDED_VARIANT_SCORERS with RNA-seq enabled

Variants are scored using AlphaGenome’s `score_variant` method, which computes effect sizes by comparing predictions between reference and alternate alleles at the specified genomic position.

#### S1.3.2 Borzoi Variant Scoring

Borzoi [35] is a sequence-to-function model that predicts genome-wide molecular phenotypes, including RNA expression levels across multiple tissues, from DNA sequence. We employed Borzoi to generate tissue-specific RNA expression predictions for genetic variants.

##### Prediction Pipeline

The Borzoi scoring pipeline consists of the following components:

1. **Variant Quality Control:** Borzoi requires high-quality SNV inputs. We apply multiple filtering steps:
  - Remove insertions and deletions (indels) by setting `max_del_len=0 and max_insert_len=0`
  - Exclude variants with non-standard bases (requiring A, C, G, T only)
  - Restrict analysis to autosomal chromosomes
  - Filter variants in ENCODE blacklist regions (100 bp window) that represent unmappable or problematic genomic regions
2. **Model Loading:** We use the pre-trained Borzoi model (`human_rep0`) from the grelu [32] resources library, which was trained on human genomic data and GTEx expression profiles.
3. **Tissue Mapping:** Similar to AlphaGenome, tissue identifiers are mapped to Borzoi’s internal GTEx sample representation using a tissue-to-sample mapping dictionary. This ensures predictions correspond to the correct biological context.
4. **Gene-Level Aggregation:** Borzoi predicts RNA levels across the entire genomic region. To obtain gene-specific predictions, the model uses gene annotations from GENCODE exon tables. The `var_to_rna` method aggregates predictions across exonic regions for each gene.
5. **Effect Size Computation:** For each variant, Borzoi generates predictions for both the reference and alternate alleles. The variant effect score is computed as the difference (or log-ratio) between alternate and reference predictions, providing a measure of expression change induced by the variant.

#### S1.3.3 Model Configuration

Key parameters for Borzoi predictions:

- Input sequence length: Model-specific training sequence length (524,288 bp)
- Reference genome: hg38
- Device: CUDA (GPU acceleration)
- Gene annotations: GENCODE exon tables for accurate gene-level aggregation

These baseline models provide strong benchmarks for evaluating our VariantFormer model’s variant effect prediction capabilities on eQTL datasets.

### S1.4 Whole Genome Variant Calling for ENCODE Cell Lines

#### Sample Data and Preprocessing

We analyzed whole genome sequencing data from six ENCODE cell lines representing diverse cancer types: A549 (lung adenocarcinoma), GM23248 (lymphoblastoid cell line), HepG2 (hepatocellular carcinoma), K562 (chronic myelogenous leukemia), NCI-H460 (lung large cell carcinoma), and Panc1 (pancreatic adenocarcinoma). Raw sequencing data were obtained from the ENCODE Data Coordination Center and processed using standardized quality control procedures. Quality assessment was performed using FastQC v0.11.9 [3] to evaluate read quality, adapter content, and sequence duplication levels. Reads were aligned to the GRCh38 reference genome (GCA_000001405.15) using BWA-MEM v0.7.17 (H. Li and Durbin 2009). Post-alignment processing included duplicate marking using Picard MarkDuplicates v2.27.5 (Broad Institute 2019) and quality score recalibration.

#### Dual-Caller Variant Detection Strategy

We implemented a variant calling approach combining germline and somatic variant detection to capture the full spectrum of genetic variation in cancer cell lines. This dual-caller strategy addresses the unique characteristics of immortalized cell lines, which may harbor both inherited germline variants and acquired somatic mutations.

#### DeepVariant Germline Calling

Germline variants were identified using DeepVariant v1.6.0 (Poplin et al. 2018) with the WGS model trained on whole genome sequencing data. DeepVariant employs a deep neural network architecture to convert genomic regions into image-like representations, enabling accurate variant calling through computer vision techniques. The pipeline was configured with 64 parallel shards distributed across make_examples, call_variants, and postprocess_variants stages to optimize computational efficiency while maintaining accuracy. Only variants with PASS filter status were retained, relying on DeepVariant’s internal quality assessment and filtering mechanisms. No additional depth or quality score thresholds were applied beyond the caller’s built-in filtering.

#### DeepSomatic Somatic Variant Detection

Somatic variants were detected using DeepSomatic v1.6.0 (Park et al. 2024) operating in tumor-only mode (WGS_TUMOR_ONLY model) with 64 parallel shards matching the DeepVariant configuration. DeepSomatic is specifically designed to identify somatic mutations in cancer samples by modeling the unique mutational patterns and allele frequency distributions characteristic of malignant cells. The algorithm accounts for potential contamination from normal cells and variable tumor purity across different cell line preparations. DeepSomatic processing included both somatic variant identification and germline variant labeling, allowing for comprehensive characterization of the mutational landscape. Variants were classified as either somatic (PASS filter) or germline-labeled based on algorithmic assessment of allele frequency patterns and genomic context. Only variants with PASS filter status were retained for somatic calls, relying on DeepSomatic’s internal quality assessment.

#### Variant Integration and Annotation

To create unified variant sets for each cell line, we developed a systematic integration procedure combining non-overlapping calls from both callers. DeepSomatic PASS variants were prioritized for somatic positions, while DeepVariant calls were retained for positions not identified as somatic. This approach ensures comprehensive coverage while avoiding double-counting of variants at identical genomic coordinates. All variants were annotated with origin type (germline or somatic) using custom VCF INFO fields. Sample names were standardized across datasets, and variants were assigned unique identifiers following the format CHROM_POS_RE_ALT. Integration was performed using BCFtools v1.15 (Danecek et al. 2021) with custom annotation scripts to maintain data provenance and quality metrics.

#### Quality Control and Validation

Comprehensive quality control measures were implemented throughout the pipeline to ensure high-confidence variant calls. Coverage analysis, mapping quality metrics, and read pairing statistics were assessed using standard genomics quality control tools. Variant quality was evaluated using multiple metrics including transition/transversion (Ts/Tv) ratios, heterozygote/homozygote ratios, and allele frequency distributions. Variant density and distribution patterns were analyzed across chromosomes to identify potential systematic biases or technical artifacts.

### S1.5 Genetic Ancestry Inference Pipeline

#### S1.5.1 Sample Data and Quality Control

We analyzed whole genome sequencing data from 2,330 samples across three major datasets: GTEx (n=943, 40.6%), ADNI (n=650, 28.0%), and 1000 Genomes Project reference populations (MAGE, n=731, 31.5%). Prior to analysis, we removed seven ENCODE cell line samples to eliminate potential artifacts from immortalized cell lines that could confound population structure analysis.

Quality control was performed using PLINK v1.90b6.24 [42]. We applied linkage disequilibrium (LD) pruning using a sliding window approach (50 SNP window, 5 SNP step, r^2^=0.2) to identify ancestry-informative markers while removing correlated variants. This resulted in 2,915,174 high-quality, independent SNPs from an initial set of 13.9M variants across the autosomal genome.

#### S1.5.2 Principal Component Analysis

We performed principal component analysis (PCA) using PLINK to capture population structure and genetic ancestry patterns. PCA was conducted on LD-pruned variants using spectral decomposition of the sample covariance matrix. We computed the first 20 principal components, which collectively explained 89.4% of the genetic variation. The first two components explained the majority of ancestry-related variation (PC1: 57.8%, PC2: 16.5%), consistent with global population structure patterns observed in human genomics studies [2, 33].

The PCA space was constructed using all samples simultaneously to ensure consistent projection and avoid batch effects. Eigenvalues were used to calculate the proportion of variance explained by each component, and sample coordinates were extracted for downstream ancestry classification.

#### S1.5.3 K-Nearest Neighbors Classifier

We implemented a K-nearest neighbors (KNN) approach for genetic ancestry inference, trained on 731 reference samples from the 1000 Genomes Project with known continental ancestry labels (African: AFR, n=196; European: EUR, n=142; East Asian: EAS, n=141; South Asian: SAS, n=139; Admixed American: AMR, n=113). These reference samples correspond to lymphoblastoid cell lines (MAGE dataset) that were excluded from ancestry inference to avoid circular validation.

The KNN classifier was configured with k=7 neighbors, distance weighting, and Euclidean distance metric in the 20-dimensional PC space. The choice of k=7 was optimized to balance local neighborhood sensitivity with classification stability across five major continental populations. Distance weighting was applied to give greater influence to closer neighbors in the PC space, which is particularly important for samples near population boundaries.

#### S1.5.4 Model Training and Validation

Classifier performance was evaluated using 5-fold cross-validation on the reference population. The KNN achieved 99.2% ± 0.8% accuracy on reference samples, demonstrating robust performance across all ancestry groups. Training was performed using scikit-learn v1.3.0 [40] with consistent random seeds for reproducibility.

#### S1.5.5 Statistical Analysis

All non-reference samples (GTEx and ADNI, n=1,593) were projected into the reference PC space and classified using the trained KNN model. We validated our approach by comparing inferred ancestry with self-reported ethnicity for samples with available demographic information. Concordance analysis was performed using standard categorical agreement metrics, with results stratified by dataset and ancestry group. This validation distinguishes between self-reported ethnicity (cultural/social identity) and genetically inferred ancestry (genomic evidence), providing insight into the relationship between these distinct but related concepts.

### S1.6 Variant Quantification and Analysis

#### S1.6.1 Computational Pipeline and Filtering Strategy

We analyzed whole genome sequencing data from four sources: ENCODE cancer cell lines (n=6), GTEx population samples (n=30), ADNI cohort (n=30), and MAGE 1000 Genomes subset (n=30). For population studies, samples were randomly selected from available cohorts.

Variant counting was performed using `bcftools`. VCF files were processed chromosome-wise (chr1-22). The pipeline uses `bcftools view -s SAMPLE -v snps/indels` for variant type filtering, followed by `bcftools query -f` ‘`[%GT]\n`’ for genotype extraction. Strict quality filtering excludes reference genotypes and missing calls using the pattern `grep -v -c -E` ‘`^(0[/|]0|.[./].)`’, which handles both phased genotypes (0|0,.|.) and unphased genotypes (0/0,./.) across different VCF formats.

#### S1.6.2 Somatic Mutation Burden Analysis

For ENCODE cell lines, somatic variants were defined as DeepSomatic calls with `FILTER=`“`PASS`”, while germline variants comprised DeepVariant calls with `FILTER=`“`PASS`” at positions where DeepSomatic did not call PASS variants (non-overlapping DeepVariant calls). This approach leverages DeepSomatic’s specialization for tumor-only somatic calling and DeepVariant’s high-confidence germline variant detection. Variant counts were generated using `bcftools`: somatic counts from `bcftools view -H deepsomatic_pass.vcf.gz | wc -l` and germline counts from `bcftools view -H deepvariant_unique.vcf.gz | wc -l`.

## References

[1] Tokenizers. https://huggingface.co/docs/tokenizers/en/index. xAccessed: 2025-10-20.

[2] 1000 Genomes Project Consortium, A. Auton, L. D. Brooks, R. M. Durbin, E. P. Garrison, H. M. Kang, J. O. Korbel, J. L. Marchini, S. McCarthy, G. A. McVean, and G. R. Abecasis. A global reference for human genetic variation. Nature, 526(7571):68–74, Oct. 2015.

[3] S. Andrews. FastQC a quality control tool for high throughput sequence data, 2010.

[4] Z. Avsec, V. Agarwal, D. Visentin, J. R. Ledsam, A. Grabska-Barwinska, K. R. Taylor, Y. Assael, J. Jumper, P. Kohli, and D. R. Kelley. Effective gene expression prediction from sequence by integrating long-range interactions. Nat. Methods, 18(10):1196–1203, Oct. 2021.

[5] Z. Avsec, N. Latysheva, J. Cheng, G. Novati, K. R. Taylor, T. Ward, C. Bycroft, L. Nicolaisen, E. Arvaniti, J. Pan, R. Thomas, V. Dutordoir, M. Perino, S. De, A. Karollus, A. Gayoso, T. Sargeant, A. Mottram, L. H. Wong, P. Drotár, A. Kosiorek, A. Senior, R. Tanburn, T. Applebaum, S. Basu, D. Hassabis, and P. Kohli. AlphaGenome: advancing regulatory variant effect prediction with a unified DNA sequence model. bioRxiv, page 2025.06.25.661532, June 2025.

[6] L. Barbadilla-Martínez, N. Klaassen, B. van Steensel, and J. de Ridder. Predicting gene expression from DNA sequence using deep learning models. Nat. Rev. Genet., 26(10):666–680, Oct. 2025.

[7] A. Bhattacharya, J. B. Hirbo, D. Zhou, W. Zhou, J. Zheng, M. Kanai, Global Biobank Meta-analysis Initiative, B. Pasaniuc, E. R. Gamazon, and N. J. Cox. Best practices for multi-ancestry, meta-analytic transcriptome-wide association studies: Lessons from the global biobank meta-analysis initiative. Cell Genom., 2(10):100180, Oct. 2022.

[8] G. Brixi, M. G. Durrant, J. Ku, M. Poli, G. Brockman, D. Chang, G. A. Gonzalez, S. H. King, D. B. Li, A. T. Merchant, M. Naghipourfar, E. Nguyen, C. Ricci-Tam, D. W. Romero, G. Sun, A. Taghibakshi, A. Vorontsov, B. Yang, M. Deng, L. Gorton, N. Nguyen, N. K. Wang, E. Adams, S. A. Baccus, S. Dillmann, S. Ermon, D. Guo, R. Ilango, K. Janik, A. X. Lu, R. Mehta, M. R. K. Mofrad, M. Y. Ng, J. Pannu, C. Re, J. C. Schmok, J. St. John, J. Sullivan, K. Zhu, G. Zynda, D. Balsam, P. Collison, A. B. Costa, T. Hernandez-Boussard, E. Ho, M.-Y. Liu, T. McGrath, K. Powell, D. P. Burke, H. Goodarzi, P. D. Hsu, and B. Hie. Genome modeling and design across all domains of life with evo 2. bioRxiv, page 2025.02.18.638918, Feb. 2025.

[9] G. Brixi, M. G. Durrant, J. Ku, M. Poli, G. Brockman, D. Chang, G. A. Gonzalez, S. H. King, D. B. Li, A. T. Merchant, M. Naghipourfar, E. Nguyen, C. Ricci-Tam, D. W. Romero, G. Sun, A. Taghibakshi, A. Vorontsov, B. Yang, M. Deng, L. Gorton, N. Nguyen, N.K. Wang, E. Adams, S.A. Baccus, S. Dillmann, S. Ermon, D. Guo, R. Ilango, K. Janik, A.X. Lu, R. Mehta, M.R.K. Mofrad, M.Y. Ng, J. Pannu, C. Ré, J.C. Schmok, J.S. John, J. Sullivan, K. Zhu, G. Zynda, D. Balsam, P. Collison, A.B. Costa, T. Hernandez-Boussard, E. Ho, M.-Y. Liu, T. McGrath, K. Powell, D.P. Burke, H. Goodarzi, P.D. Hsu, and B.L. Hie. Genome modeling and design across all domains of life with evo 2. Genomics, (biorxiv;2025.02.18.638918v1), Feb. 2025.

[10] M. N. Cabili, C. Trapnell, L. Goff, M. Koziol, B. Tazon-Vega, A. Regev, and J. L. Rinn. Integrative annotation of human large intergenic noncoding RNAs reveals global properties and specific subclasses. Genes Dev., 25(18):1915–1927, Sept. 2011.

[11] H. Cai, T. Zhao, Y. Pang, X. Fu, Z. Ren, S. Quan, and L. Jia. Systemic inflammatory markers in ageing, alzheimer’s disease and other dementias. Brain, 148(2):480–492, Feb. 2025.

[12] S. Carpenter, D. Aiello, M. K. Atianand, E. P. Ricci, P. Gandhi, L. L. Hall, M. Byron, B. Monks, M. Henry-Bezy, J. B. Lawrence, L. A. J. O’Neill, M. J. Moore, D. R. Caffrey, and K. A. Fitzgerald. A long noncoding RNA mediates both activation and repression of immune response genes. Science, 341(6147):789–792, Aug. 2013.

[13] H. Dalla-Torre, L. Gonzalez, J. Mendoza-Revilla, N. Lopez Carranza, A. H. Grzywaczewski, F. Oteri, C. Dallago, E. Trop, B. P. de Almeida, H. Sirelkhatim, G. Richard, M. Skwark, K. Beguir, M. Lopez, and T. Pierrot. Nucleotide transformer: building and evaluating robust foundation models for human genomics. Nat. Methods, 22(2):287–297, Feb. 2025.

[14] P. Danecek, J. K. Bonfield, J. Liddle, J. Marshall, V. Ohan, M. O. Pollard, A. Whitwham, T. Keane, S. A. McCarthy, R. M. Davies, and H. Li. Twelve years of SAMtools and BCFtools. Gigascience, 10(2):giab008, Feb. 2021.

[15] T. Dao, D. Y. Fu, S. Ermon, A. Rudra, and C. Ré. FlashAttention: Fast and memory-efficient exact attention with IO-awareness. arXiv [cs.LG], May 2022.

[16] C. A. de Leeuw, J. M. Mooij, T. Heskes, and D. Posthuma. MAGMA: generalized gene-set analysis of GWAS data. PLoS Comput. Biol., 11(4):e1004219, Apr. 2015.

[17] N. Ehsan, B. M. Kotis, S. E. Castel, E. J. Song, N. Mancuso, and P. Mohammadi. Haplotype-aware modeling of cis-regulatory effects highlights the gaps remaining in eQTL data. Nat. Commun., 15(1):522, Jan. 2024.

[18] ENCODE Project Consortium et al. Expanded encyclopaedias of DNA elements in the human and mouse genomes. Nature, 583(7818):699–710, July 2020.

[19] J. I. Fuxman Bass. Understanding the logic and grammar of cis-regulatory elements. Nat. Rev. Genet., 26(10):665, Oct. 2025.

[20] E. R. Gamazon, H. E. Wheeler, K. P. Shah, S. V. Mozaffari, K. Aquino-Michaels, R. J. Carroll, A. E. Eyler, J. C. Denny, GTEx Consortium, D. L. Nicolae, N. J. Cox, and H. K. Im. A gene-based association method for mapping traits using reference transcriptome data. Nat. Genet., 47(9):1091–1098, Sept. 2015.

[21] GTEx Consortium. The GTEx consortium atlas of genetic regulatory effects across human tissues. Science, 369(6509):1318–1330, Sept. 2020.

[22] A. Gusev, A. Ko, H. Shi, G. Bhatia, W. Chung, B. W. J. H. Penninx, R. Jansen, E. J. C. de Geus, D. I. Boomsma, F. A. Wright, P. F. Sullivan, E. Nikkola, M. Alvarez, M. Civelek, A. J. Lusis, T. Lehtimäki, E. Raitoharju, M. Kähönen, I. Seppälä, O. T. Raitakari, J. Kuusisto, M. Laakso, A. L. Price, P. Pajukanta, and B. Pasaniuc. Integrative approaches for large-scale transcriptome-wide association studies. Nat. Genet., 48(3):245–252, Mar. 2016.

[23] C. Huang, R. W. Shuai, P. Baokar, R. Chung, R. Rastogi, P. Kathail, and N. M. Ioannidis. Personal transcriptome variation is poorly explained by current genomic deep learning models. Nat. Genet., 55(12):2056–2059, Dec. 2023.

[24] H. I. L. Jacobs, D. A. Hopkins, H. C. Mayrhofer, E. Bruner, F. W. van Leeuwen, W. Raaijmakers, and J. D. Schmahmann. The cerebellum in alzheimer’s disease: evaluating its role in cognitive decline. Brain, 141(1):37–47, Jan. 2018.

[25] Y. Ji, Z. Zhou, H. Liu, and R. V. Davuluri. DNABERT: pre-trained bidirectional encoder representations from transformers model for DNA-language in genome. Bioinformatics, 37(15):2112–2120, Aug. 2021.

[26] G. A. Jindal and E. K. Farley. Enhancer grammar in development, evolution, and disease: dependencies and interplay. Dev. Cell, 56(5):575–587, Mar. 2021.

[27] J. K. Johnson, E. Head, R. Kim, A. Starr, and C. W. Cotman. Clinical and pathological evidence for a frontal variant of alzheimer disease. Arch. Neurol., 56(10):1233–1239, Oct. 1999.

[28] D. R. Kelley, J. Snoek, and J. L. Rinn. Basset: learning the regulatory code of the accessible genome with deep convolutional neural networks. Genome Res., 26(7):990–999, July 2016.

[29] N. Kerimov, J. D. Hayhurst, K. Peikova, J. R. Manning, P. Walter, L. Kolberg, M. Samoviča, M. P. Sakthivel, I. Kuzmin, S. J. Trevanion, T. Burdett, S. Jupp, H. Parkinson, I. Papatheodorou, A. D. Yates, D. R. Zerbino, and K. Alasoo. A compendium of uniformly processed human gene expression and splicing quantitative trait loci. Nat. Genet., 53(9):1290–1299, Sept. 2021.

[30] K. L. Keys, A. C. Y. Mak, M. J. White, W. L. Eckalbar, A. W. Dahl, J. Mefford, A. V. Mikhaylova, M. G. Contreras, J. R. Elhawary, C. Eng, D. Hu, S. Huntsman, S. S. Oh, S. Salazar, M. A. Lenoir, J. C. Ye, T. A. Thornton, N. Zaitlen, E. G. Burchard, and C. R. Gignoux. On the cross-population generalizability of gene expression prediction models. PLoS Genet., 16(8):e1008927, Aug. 2020.

[31] L. Kozma and J. Voderholzer. Theoretical analysis of byte-pair encoding. arXiv [cs.DS], Nov. 2024.

[32] A. Lal, L. Gunsalus, S. Nair, T. Biancalani, and G. Eraslan. gReLU: a comprehensive framework for DNA sequence modeling and design. Nat. Methods, pages 1–5, Oct. 2025.

[33] J. Z. Li, D. M. Absher, H. Tang, A. M. Southwick, A. M. Casto, S. Ramachandran, H. M. Cann, G. S. Barsh, M. Feldman, L. L. Cavalli-Sforza, and R. M. Myers. Worldwide human relationships inferred from genome-wide patterns of variation. Science, 319(5866):1100–1104, Feb. 2008.

[34] W.-W. Liao, M. Asri, J. Ebler, D. Doerr, M. Haukness, G. Hickey, S. Lu, J. K. Lucas, J. Monlong, H. J. Abel, S. Buonaiuto, X. H. Chang, H. Cheng, J. Chu, V. Colonna, J. M. Eizenga, X. Feng, C. Fischer, R. S. Fulton, S. Garg, C. Groza, A. Guarracino, W. T. Harvey, S. Heumos, K. Howe, M. Jain, T.-Y. Lu, C. Markello, F. J. Martin, M. W. Mitchell, K. M. Munson, M. N. Mwaniki, A. M. Novak, H. E. Olsen, T. Pesout, D. Porubsky, P. Prins, J. A. Sibbesen, J. Sirén, C. Tomlinson, F. Villani, M. R. Vollger, L. L. Antonacci-Fulton, G. Baid, C. A. Baker, A. Belyaeva, K. Billis, A. Carroll, P.-C. Chang, S. Cody, D. E. Cook, R. M. Cook-Deegan, O. E. Cornejo, M. Diekhans, P. Ebert, S. Fairley, O. Fedrigo, A. L. Felsenfeld, G. Formenti, A. Frankish, Y. Gao, N. A. Garrison, C. G. Giron, R. E. Green, L. Haggerty, K. Hoekzema, T. Hourlier, H. P. Ji, E. E. Kenny, B. A. Koenig, A. Kolesnikov, J. O. Korbel, J. Kordosky, S. Koren, H. Lee, A. P. Lewis, H. Magalhães, S. Marco-Sola, P. Marijon, A. McCartney, J. McDaniel, J. Mountcastle, M. Nattestad, S. Nurk, N. D. Olson, A. B. Popejoy, D. Puiu, M. Rautiainen, A. A. Regier, A. Rhie, S. Sacco, A. D. Sanders, V. A. Schneider, B. I. Schultz, K. Shafin, M. W. Smith, H. J. Sofia, A. N. Abou Tayoun, F. Thibaud-Nissen, F. F. Tricomi, J. Wagner, B. Walenz, J. M. D. Wood, A. V. Zimin, G. Bourque, M. J. P. Chaisson, P. Flicek, A. M. Phillippy, J. M. Zook, E. E. Eichler, D. Haussler, T. Wang, E. D. Jarvis, K. H. Miga, E. Garrison, T. Marschall, I. M. Hall, H. Li, and B. Paten. A draft human pangenome reference. Nature, 617(7960):312–324, May 2023.

[35] J. Linder, D. Srivastava, H. Yuan, V. Agarwal, and D. R. Kelley. Predicting RNA-seq coverage from DNA sequence as a unifying model of gene regulation. Nat. Genet., 57(4):949–961, Apr. 2025.

[36] A. V. Mikhaylova and T. A. Thornton. Accuracy of gene expression prediction from genotype data with PrediXcan varies across and within continental populations. Front. Genet., 10(261):261, Apr. 2019.

[37] J. M. Mudge et al. GENCODE 2025: reference gene annotation for human and mouse. Nucleic Acids Res., 53(D1):D966–D975, Jan. 2025.

[38] E. Nguyen, M. Poli, M. Faizi, A. Thomas, C. Birch-Sykes, M. Wornow, A. Patel, C. Rabideau, S. Massaroli, Y. Bengio, S. Ermon, S. A. Baccus, and C. Ré. HyenaDNA: Long-range genomic sequence modeling at single nucleotide resolution. arXiv [cs.LG], June 2023.

[39] Ofir Press, N. A. Smith, and M. Lewis. Train short, test long: Attention with linear biases enables input length extrapolation. arXiv [cs.CL], Aug. 2021.

[40] F. Pedregosa, G. Varoquaux, A. Gramfort, V. Michel, B. Thirion, O. Grisel, M. Blondel, G. Louppe, P. Prettenhofer, R. Weiss, R. J. Weiss, J. Vanderplas, A. Passos, D. Cournapeau, M. Brucher, M. Perrot, and E. Duchesnay. Scikit-learn: Machine learning in python. J. Mach. Learn. Res., 12:2825–2830, Feb. 2011.

[41] R. C. Petersen, P. S. Aisen, L. A. Beckett, M. C. Donohue, A. C. Gamst, D. J. Harvey, C. R. Jack, Jr, W. J. Jagust, L. M. Shaw, A. W. Toga, J. Q. Trojanowski, and M. W. Weiner. Alzheimer’s disease neuroimaging initiative (ADNI): clinical characterization. Neurology, 74(3):201–209, Jan. 2010.

[42] S. Purcell, B. Neale, K. Todd-Brown, L. Thomas, M. A. R. Ferreira, D. Bender, J. Maller, P. Sklar, P. I. W. de Bakker, M. J. Daly, and P. C. Sham. PLINK: a tool set for whole-genome association and population-based linkage analyses. Am. J. Hum. Genet., 81(3):559–575, Sept. 2007.

[43] M. H. Quiver and J. Lachance. Adaptive eQTLs reveal the evolutionary impacts of pleiotropy and tissue-specificity while contributing to health and disease. HGG Adv., 3(1):100083, Jan. 2022.

[44] Roadmap Epigenomics Consortium et al. Integrative analysis of 111 reference human epigenomes. Nature, 518(7539):317–330, Feb. 2015.

[45] A. D. Roses, M. W. Lutz, H. Amrine-Madsen, A. M. Saunders, D. G. Crenshaw, S. S. Sundseth, M. J. Huentelman, K. A. Welsh-Bohmer, and E. M. Reiman. A TOMM40 variable-length polymorphism predicts the age of late-onset alzheimer’s disease. Pharmacogenomics J., 10(5):375–384, Oct. 2010.

[46] A. Saeed, O. Lopez, A. Cohen, and S. E. Reis. Cardiovascular disease and alzheimer’s disease: The heart-brain axis. J. Am. Heart Assoc., 12(21):e030780, Nov. 2023.

[47] A. Serrano-Pozo, S. Das, and B. T. Hyman. APOE and alzheimer’s disease: advances in genetics, pathophysiology, and therapeutic approaches. Lancet Neurol., 20(1):68–80, Jan. 2021.

[48] A. E. Spiro, X. Tu, Y. Sheng, A. Sasse, R. Hosseini, M. Chikina, and S. Mostafavi. A scalable approach to investigating sequence-to-function predictions from personal genomes. bioRxiv, page 2025.02.21.639494, Feb. 2025.

[49] D. J. Taylor, S. B. Chhetri, M. G. Tassia, A. Biddanda, S. M. Yan, G. L. Wojcik, A. Battle, and R. C. McCoy. Sources of gene expression variation in a globally diverse human cohort. Nature, 632(8023):122–130, Aug. 2024.

[50] I. Ulitsky and D. P. Bartel. lincRNAs: genomics, evolution, and mechanisms. Cell, 154(1):26–46, July 2013.

[51] I. Virshup, S. Rybakov, F. J. Theis, P. Angerer, and F. A. Wolf. anndata: Annotated data. bioRxiv, page 2021.12.16.473007, Dec. 2021.

[52] M. Wainberg, N. Sinnott-Armstrong, N. Mancuso, A. N. Barbeira, D. A. Knowles, D. Golan, R. Ermel, A. Ruusalepp, T. Quertermous, K. Hao, J. L. M. Björkegren, H. K. Im, B. Pasaniuc, M. A. Rivas, and A. Kundaje. Opportunities and challenges for transcriptome-wide association studies. Nat. Genet., 51(4):592–599, Apr. 2019.

[53] H. Wang, D. Lou, and Z. Wang. Crosstalk of genetic variants, allele-specific DNA methylation, and environmental factors for complex disease risk. Front. Genet., 9:695, 2018.

[54] X. Wang, F. Li, Y. Zhang, S. Imoto, H.-H. Shen, S. Li, Y. Guo, J. Yang, and J. Song. Deep learning approaches for non-coding genetic variant effect prediction: current progress and future prospects. Brief. Bioinform., 25(5):bbae446, July 2024.

[55] M. W. Weiner, P. S. Aisen, C. R. Jack, Jr, W. J. Jagust, J. Q. Trojanowski, L. Shaw, A. J. Saykin, J. C. Morris, N. Cairns, L. A. Beckett, A. Toga, R. Green, S. Walter, H. Soares, P. Snyder, E. Siemers, W. Potter, P. E. Cole, M. Schmidt, and Alzheimer’s Disease Neuroimaging Initiative. The alzheimer’s disease neuroimaging initiative: progress report and future plans. Alzheimers. Dement., 6(3):202–11.e7, May 2010.

[56] X. Wen, F. Luca, and R. Pique-Regi. Cross-population joint analysis of eQTLs: fine mapping and functional annotation. PLoS Genet., 11(4):e1005176, Apr. 2015.

[57] Q. Yuan, X. Liang, C. Xue, W. Qi, S. Chen, Y. Song, H. Wu, X. Zhang, C. Xiao, and J. Chen. Altered anterior cingulate cortex subregional connectivity associated with cognitions for distinguishing the spectrum of pre-clinical alzheimer’s disease. Front. Aging Neurosci., 14:1035746, Dec. 2022.

[58] J. Zhang and H. Zhao. eQTL studies: from bulk tissues to single cells. J. Genet. Genomics, 50(12):925–933, Dec. 2023.

[59] H. Zhao, Z. Sun, J. Wang, H. Huang, J.-P. Kocher, and L. Wang. CrossMap: a versatile tool for coordinate conversion between genome assemblies. Bioinformatics, 30(7):1006–1007, Apr. 2014.

[60] J. Zhou and O. G. Troyanskaya. Predicting effects of noncoding variants with deep learning-based sequence model. Nat. Methods, 12(10):931–934, Oct. 2015.

[61] S. Zunjarrao and M. C. Gambetta. Principles of long-range gene regulation. Curr. Opin. Genet. Dev., 91(102323):102323, Apr. 2025.

